# Molecular determinants of ligand recognition and efficacy in the TRPM3 voltage sensor-like domain

**DOI:** 10.64898/2026.09.27.753811

**Authors:** Vikram Dalal, Vipin A. Rangari, Samuel Eriksson Lidbrink, K. Trent Moreland, Erik Lindahl, Rebecca J. Howard, Susruta Majumdar, Wayland W. L. Cheng

**Affiliations:** Department of Anesthesiology, Washington University School of Medicine, St. Louis, MO, USA; Center for Clinical Pharmacology and Washington University Pain Center, Washington University School of Medicine, St. Louis, MO, USA; Department of Biochemistry and Biophysics, SciLifeLab, Stockholm University, Solna, Sweden; Department of Applied Physics, SciLifeLab, KTH Royal Institute of Technology, Solna, Sweden; Department of Physics, Chemistry and Biology, Linköping University, Linköping, Sweden; Department of Chemistry, University of Illinois at Urbana-Champaign, Urbana, IL, USA

## Abstract

Transient receptor potential melastatin 3 (TRPM3) is a calcium-permeable ion channel implicated in chronic pain and developmental and epileptic encephalopathies. The voltage sensor-like domain (VSLD) is a major site of pharmacological modulation, but the molecular determinants of ligand recognition and efficacy remain incompletely understood. Here we combine cryo-electron microscopy, molecular dynamics simulations, mutagenesis, and medicinal chemistry to investigate the interactions of TRPM3 with the inhibitors isosakuranetin, maprotiline, and diclofenac. We show that structurally diverse ligands occupy a common VSLD cavity but engage distinct interaction networks. Functional and computational analyses identify ligand-specific residue dependencies and support a halogen bond between diclofenac and D947 that contributes to inhibitor potency. Furthermore, esterification or amidation of the diclofenac carboxylate converts inhibition into potentiation, and mutagenesis and molecular dynamics simulations support partially overlapping VSLD configurations associated with these opposing effects. Together, these findings establish the TRPM3 VSLD as an adaptable allosteric pocket in which ligand-specific interactions shape recognition, potency, and efficacy.

## Introduction

Transient receptor potential melastatin 3 (TRPM3) is a non-selective cation channel expressed in sensory neurons and other excitable tissues where it contributes to thermosensation^1^, synaptic transmission^2^, and vascular physiology^3^. Genetic and pharmacological studies have established TRPM3 as a key mediator of inflammatory and neuropathic pain^4–7^, while human gain-of-function mutations in TRPM3 cause developmental and epileptic encephalopathies characterized by seizures, intellectual disability, and neurodevelopmental deficits^8, 9^. Accordingly, inhibition of TRPM3 has emerged as a promising therapeutic strategy for chronic pain^5, 10–12^ and TRPM3-associated channelopathies^13^.

Recent cryo-EM studies have advanced our understanding of TRPM3 gating and pharmacology^14^. Structures of TRPM3 bound to the agonist CIM0216, as well as the inhibitors primidone and isosakuranetin, identified the voltage sensor-like domain (VSLD) as a major site for chemical modulation of channel activity^15–17^. CIM0216 and these inhibitors occupy overlapping regions of the VSLD yet are associated with distinct conformational states of the channel, establishing this pocket as a key allosteric site modulating TRPM3 function. More recently, mutational and structural analyses demonstrated that subtle changes in channel sequence combined with ligand chemistry can alter the direction of channel modulation, highlighting an unexpected degree of pharmacological plasticity within the VSLD^16^. Despite the identification of the VSLD as a common binding site for TRPM3 modulators, the molecular basis of inhibitor recognition remains incompletely understood. Structural information for TRPM3 inhibitors such as maprotiline and diclofenac is lacking, limiting understanding of how specific interactions contribute to ligand potency. Moreover, whether subtle changes in ligand chemistry alone can switch the direction of TRPM3 modulation, and what molecular features distinguish inhibition from potentiation within the VSLD, remain unclear. Throughout this study, we use the term inhibitor rather than antagonist to describe compounds that suppress TRPM3 activity because VSLD-binding ligands also inhibit channel activation by pregnenolone sulfate (PS), which acts through distinct binding sites^17^.

Here, we combine cryo-EM, molecular dynamics simulations, mutagenesis, and medicinal chemistry to elucidate molecular principles governing ligand recognition at the TRPM3 VSLD. We identify distinct interaction fingerprints for multiple inhibitor classes, provide evidence that halogen bonding contributes to diclofenac potency, and demonstrate that subtle modification of diclofenac alone can switch its effect on WT TRPM3 from inhibition to potentiation. Functional and computational analyses further support distinct, partially overlapping modes of VSLD engagement associated with these opposing effects.

## Results

### Structurally diverse TRPM3 inhibitors converge on a common VSLD pocket

To define molecular determinants of inhibitor recognition in TRPM3, we determined cryo-EM structures of the channel bound to the inhibitors isosakuranetin, maprotiline, and diclofenac. Despite substantial differences in their chemical scaffolds, all three ligands occupy the voltage sensor-like domain (VSLD) cavity previously shown to bind the agonist CIM0216 and the inhibitor primidone^16, 17^, establishing this region as a common recognition site for structurally diverse TRPM3 modulators. Notably, isosakuranetin, maprotiline, and diclofenac are neutral, cationic, and anionic ligands, respectively, underscoring the chemical diversity accommodated by the VSLD.

Prior to structure determination, we confirmed inhibition of mouse TRPM3 using a calcium mobilization assay with responses elicited by 50 μM pregnenolone sulfate (PS). Maprotiline and diclofenac inhibited TRPM3 with IC_50_ values of 1.8 μM and 20 μM, respectively, consistent with previous reports^10, 11, 18^. Isosakuranetin produced a biphasic concentration-response relationship that was best fit by two inhibitory components with IC_50_ values of 0.1 μM and 6.5 μM (Fig. 1A). A recent study demonstrated that the R-enantiomer of isosakuranetin is more potent than the S-enantiomer^16^, raising the possibility that the two inhibitory components reflect differential contributions from the two stereoisomers.

**Figure 1:**
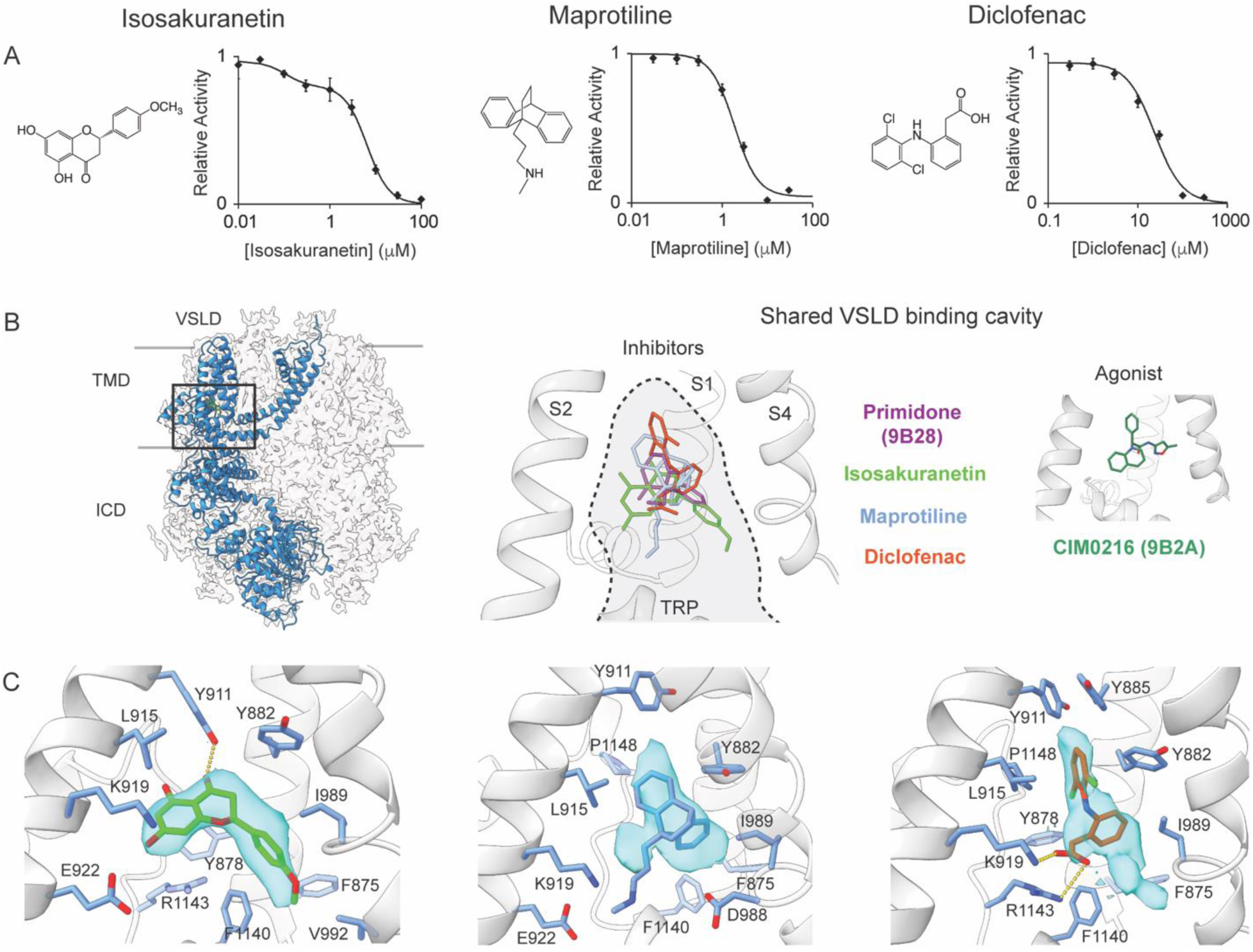
Structurally diverse TRPM3 inhibitors occupy a shared VSLD binding cavity while engaging distinct interaction networks. **(A)** Chemical structures and concentration–response relationships for inhibition of mouse TRPM3 by isosakuranetin, maprotiline, and diclofenac. TRPM3 activity was measured using a calcium mobilization assay in response to 50 μM pregnenolone sulfate (PS). Data are mean ± SEM (n = 8–25). Isosakuranetin inhibition was best described by a biphasic concentration–response relationship. **(B)** Location of the voltage sensor-like domain (VSLD) binding cavity within TRPM3. Left: overview of the TRPM3 structure with the VSLD highlighted. Center: overlay of inhibitor-bound structures showing that primidone (PDB 9B28), isosakuranetin, maprotiline, and diclofenac occupy a shared cavity within the VSLD. Right: the agonist CIM0216 (PDB 9B2A) binds an overlapping region of the same pocket. **(C)** Cryo-EM densities and binding poses of isosakuranetin, maprotiline, and diclofenac within the VSLD cavity. Ligands are shown as sticks with corresponding cryo-EM densities. Side chains of residues within 4 Å of each ligand are shown as sticks. Dashed lines indicate putative hydrogen bond or ionic interactions in the cryo-EM models.

Structures were then determined using a previously described mouse TRPM3 construct lacking the distal C-terminal region (residues 1–1344)^14^. Cryo-EM reconstructions were obtained at overall resolutions of 3.6 Å for isosakuranetin, 3.4 Å for maprotiline, and 3.2 Å for diclofenac (Supplementary Figs. 1–3, Table 1). All three structures adopt a closed-state conformation closely resembling previously reported apo and inhibitor-bound structures, with constrictions at G1056 in the selectivity filter and N1115 at the intracellular end of S6 (Supplementary Fig. 4A). To preserve the disulfide bond between C1067 and C1084 in the pore loop, which contributes to inhibition by isosakuranetin^19^, structures were determined in the absence of the reducing agent DTT. Consistent with this expectation, density corresponding to the disulfide bond was observed in all three structures and was most evident in the isosakuranetin-bound reconstruction (Supplementary Fig. 4B). Samples were also prepared in the presence of 200 μM diC8-PIP_2_, which is required for TRPM3 activity, although a corresponding diC8-PIP_2_ density could not be unambiguously assigned.

Non-protein density corresponding to each inhibitor was observed within the VSLD cavity encompassed by the S1–S4 helices (Fig. 1C, Supplementary Fig. 2 and 3). The density for isosakuranetin permitted unambiguous modeling of the ligand and was best fit by the R-enantiomer, consistent with the greater inhibitory potency of this stereoisomer^16^. In contrast, the densities for maprotiline and diclofenac were less well defined, particularly for the maprotiline methylpropanamine side chain and the overall orientation of diclofenac. For diclofenac, the density was consistent with two possible binding configurations that differed in the orientation of the 1,3-dichlorophenyl ring (Supplementary Fig 4C). These observations suggest configurational heterogeneity within the VSLD and motivated further analysis using molecular dynamics simulations. Nevertheless, overlay of primidone (PDB 9B28), isosakuranetin, maprotiline, and diclofenac demonstrates that chemically diverse inhibitors occupy a common cavity within the VSLD, while the agonist CIM0216 (PDB 9B2A) binds an overlapping region of the same pocket (Fig. 1B).

Although all three inhibitors occupy the same VSLD cavity, inspection of the structures suggests distinct interaction networks (Fig. 1C). Isosakuranetin forms a hydrogen bond interaction with Y911, and hydrophobic contacts involving F1140, I989, L915, and aromatic residues lining the pocket. In contrast, maprotiline and diclofenac adopt positions shifted toward the extracellular side of the VSLD and engage different subsets of residues. The N-methylpropylamine substituent of maprotiline is near E922 and D988, while the carboxylate group on the phenylacetic ring of diclofenac forms salt bridge interactions with K919 and R1143. These observations indicate that a common VSLD pocket accommodates chemically diverse inhibitors with varying charge through distinct molecular interactions.

### Molecular determinants of maprotiline inhibition

To define the molecular interactions that mediate inhibition, we combined molecular dynamics (MD) simulations with functional analysis of VSLD mutants. Isosakuranetin was recently subjected to mutational analysis^16^, prompting us to focus on the structurally distinct inhibitors maprotiline and diclofenac. MD simulations were initiated using the cryo-EM structure to examine the stability and conformational landscape of maprotiline within the VSLD pocket. Four independent 500-ns simulations were performed with maprotiline bound to each VSLD site, yielding 16 trajectories (Supplementary Table 2). Maprotiline remained within the VSLD but deviated from its cryo-EM position by ∼4–5 Å RMSD (Supplementary Fig. 5). Clustering identified four pose ensembles, with two predominant clusters accounting for 72% of sampled conformations. In these poses, maprotiline shifts toward S4 and the intracellular VSLD, where its tetracyclic scaffold forms hydrophobic contacts with F1140, I989, Y882, Y878, and F875 and its protonated amine forms a salt bridge with D988 (Fig. 2A–B, Supplementary Fig. 5). By contrast, the two less populated clusters more closely resemble the cryo-EM pose, retaining the position of the methylpropanamine while exhibiting a tilt of the tetracyclic rings toward S2 that promotes interactions with L915.

**Figure 2:**
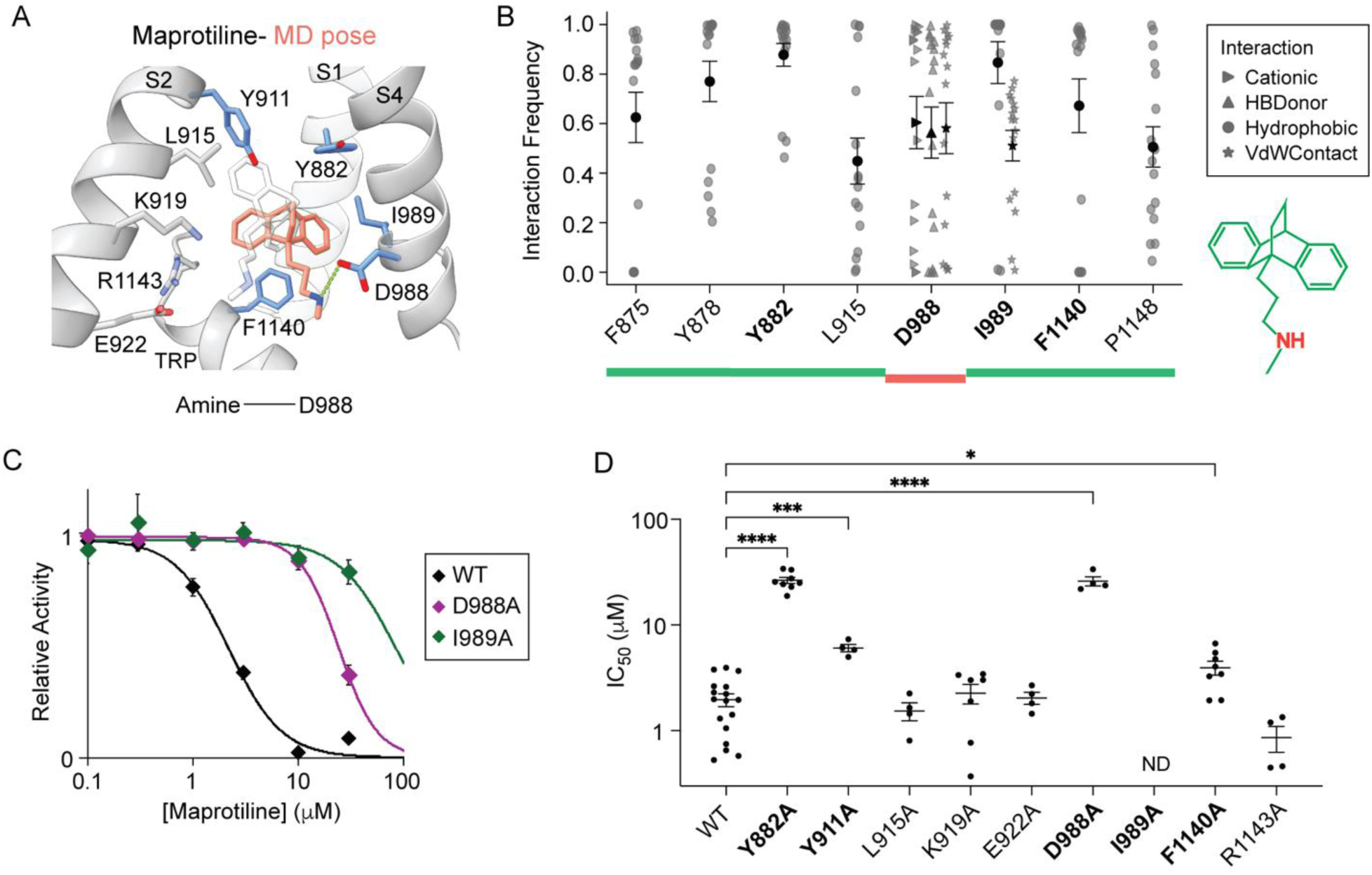
Molecular determinants of maprotiline recognition within the TRPM3 VSLD. **(A)** Representative binding pose of maprotiline derived from MD simulations initiated from the cryo-EM structure. The protonated amine is positioned near D988 forming an ionic interaction. Residues within 4 Å of the ligand are shown as sticks (blue-mutations decrease potency). The semi-transparent gray ligand indicates the starting cryo-EM pose. **(B)** Interaction frequencies between maprotiline and VSLD residues from the MD simulations shown for individual replicates with mean ± SEM. Residues highlighted in bold exhibited significant effects on maprotiline inhibition in mutagenesis experiments. The maprotiline scaffold beneath the plot highlights the two major pharmacophoric regions evaluated in the simulations. Colored bars indicate residues that primarily interact with the tetracyclic scaffold (green) or the protonated amine (red) of maprotiline. **(C)** Representative concentration–response relationships for inhibition of PS-evoked TRPM3 activity by maprotiline in WT and mutants. TRPM3 activity was measured using a calcium mobilization assay in response to PS. Data are shown as mean ± SEM. **(D)** Summary of maprotiline IC_50_ values for WT and mutants showing mean ± SEM. Statistical significance was determined by one-way ANOVA with Dunnett’s multiple-comparisons test relative to WT. ND-minimal inhibition within the concentration range tested. * p<0.05, ** p<0.01, *** p<0.001, **** p<0.0001.

**Figure 3:**
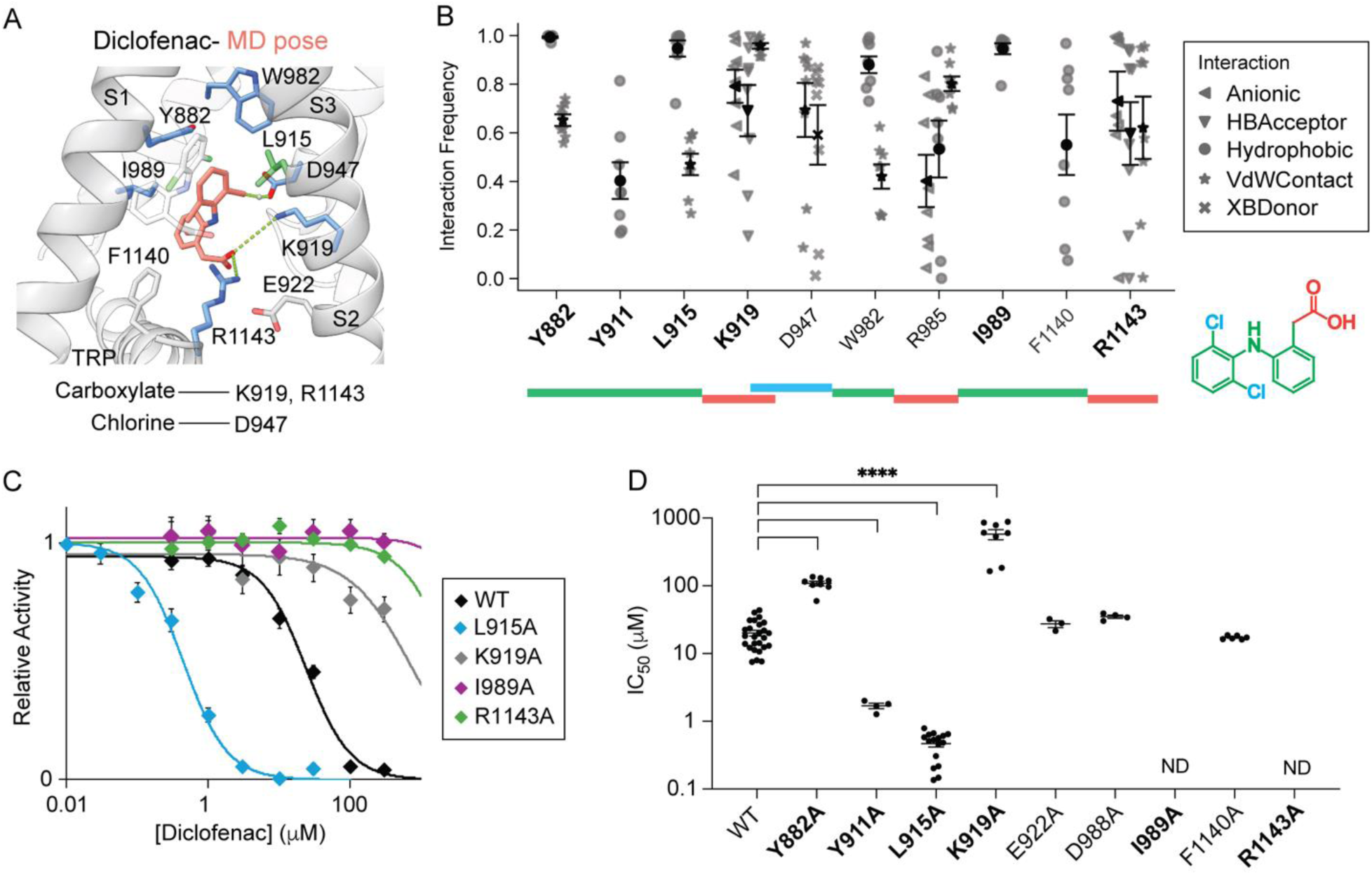
Molecular determinants of diclofenac recognition within the TRPM3 VSLD. **(A)** Representative binding pose of diclofenac derived from MD simulations initiated from configuration 1 of the cryo-EM structure. The carboxylate group forms ionic interactions with K919 and R1143, whereas a chlorine is positioned near D947. Residues within 4 Å of the ligand are shown as sticks (blue-mutations decrease potency; green-mutations increase potency). The semi-transparent gray ligand indicates the starting cryo-EM pose. **(B)** Interaction frequencies between diclofenac and VSLD residues from MD simulations shown for individual replicates with mean ± SEM. Residues highlighted in bold exhibited significant effects on diclofenac inhibition in mutagenesis experiments. The diclofenac scaffold beneath the plot highlights the three major pharmacophoric regions evaluated in the simulations. Colored bars indicate residues that primarily interact with the aromatic scaffold (green), chlorines (cyan) or the carboxylate group (red) of diclofenac. **(C)** Representative concentration–response relationships for inhibition of PS-evoked TRPM3 activity by diclofenac in WT and mutants. TRPM3 activity was measured using a calcium mobilization assay in response to PS. Data are shown as mean ± SEM. **(D)** Summary of diclofenac IC_50_ values for WT and mutants showing mean ± SEM. Statistical significance was determined by one-way ANOVA with Dunnett’s multiple-comparisons test relative to WT. ND-minimal inhibition within the concentration range tested. ****p < 0.0001.

To experimentally test the interactions predicted by the simulations, we quantified maprotiline inhibition of alanine mutants using calcium mobilization assays. We first determined PS concentration response relationships for VSLD mutants. Y882A, I989A, and F1140A exhibited substantially increased PS sensitivity relative to WT (Supplementary Fig. 6). We therefore measured inhibition at PS concentrations producing approximately EC_50-70_ activation: 3 μM for Y882A and I989A, 5 μM for F1140A, and 50 μM for WT and the remaining mutants. Mutations predicted to disrupt interactions in the dominant MD-derived poses produced the largest effects on inhibition. F1140A, I989A, Y911A, Y882A, and D988A significantly reduced or abolished maprotiline inhibition (Fig. 2C and 2D, Supplementary Fig. 7). In contrast, residues located near maprotiline in the cryo-EM structure but not predicted to form contacts in the simulations, including L915A, K919A, E922A, and R1143A, retained near WT sensitivity. No measurable TRPM3 activity was detected for Y878A, F875A or P1148A, precluding evaluation of their contributions to maprotiline inhibition. Together, mutagenesis and MD define a maprotiline interaction fingerprint comprising a D988 salt bridge and hydrophobic contacts with I989, Y882, Y911, and F1140.

### Molecular determinants of diclofenac inhibition

The cryo-EM density for diclofenac was consistent with two possible binding configurations. In configuration 1, the dichlorophenyl ring is positioned toward the extracellular side of the VSLD, whereas in configuration 2 the dichlorophenyl ring is oriented toward the intracellular side (Supplementary Fig. 4C). To evaluate these configurations, we performed two independent 500-ns MD simulations for each starting model, yielding eight trajectories per configuration (Supplementary Table 2). In simulations initiated from configuration 1, diclofenac moved toward S3 and adopted poses that deviated from the cryo-EM model by 3–6 Å RMSD (Supplementary Fig. 8). These poses frequently positioned a chlorine near D947 at an interatomic distance of ∼3 Å, with an orientation consistent with a putative halogen bond (Fig. 3A and 3B, Fig. 5B). The carboxylate group formed salt bridge interactions with K919, R985, and R1143, while the dichlorophenyl ring contacted L915, W885, and W892 and the phenylacetic ring contacted I989 and Y882. In contrast, simulations initiated from configuration 2 showed rotation of the phenylacetic ring toward S3 and formation of hydrophobic contacts with F1140 in the TRP domain that were not observed in configuration 1 (Supplementary Fig. 9).

To test these predicted interactions, we examined the effects of VSLD mutations on diclofenac inhibition. D947A did not exhibit measurable PS-evoked activity, precluding direct evaluation of the putative halogen bond. Also, R985A and W892A were similarly nonfunctional. K919A and R1143A markedly reduced diclofenac inhibition, whereas D988A and E922A had little effect, consistent with a binding mode in which the diclofenac carboxylate engages K919 and R1143 (Fig. 3C and 3D). I989A and Y882A abolished or substantially reduced inhibition, whereas F1140A had minimal effect. Notably, F1140 contacts diclofenac in the cryo-EM model and in MD-equilibrated poses derived from configuration 2, but not in the dominant poses derived from configuration 1 (Supplementary Figs. 8 and 9). The minimal effect of F1140A therefore supports poses derived from configuration 1 as candidate inhibitory configurations.

L915A increased diclofenac potency by approximately 60-fold (Fig. 3C and 3D). To investigate this effect, we performed MD simulations with the L915A mutation introduced in silico. Relative to WT, L915A showed more stable occupancy of the candidate inhibitory configuration derived from configuration 1, compared to simulations initiated from configuration 2 (Fig. 4E, Supplementary Fig. 10 and 11). Collectively, the mutagenesis and MD simulations define a diclofenac interaction fingerprint characterized by ionic interactions with K919 and R1143, hydrophobic contacts with Y882 and I989, and a putative halogen bond with D947. Furthermore, the combined data define a candidate binding configuration derived from configuration 1 that contributes to diclofenac inhibition.

**Figure 4:**
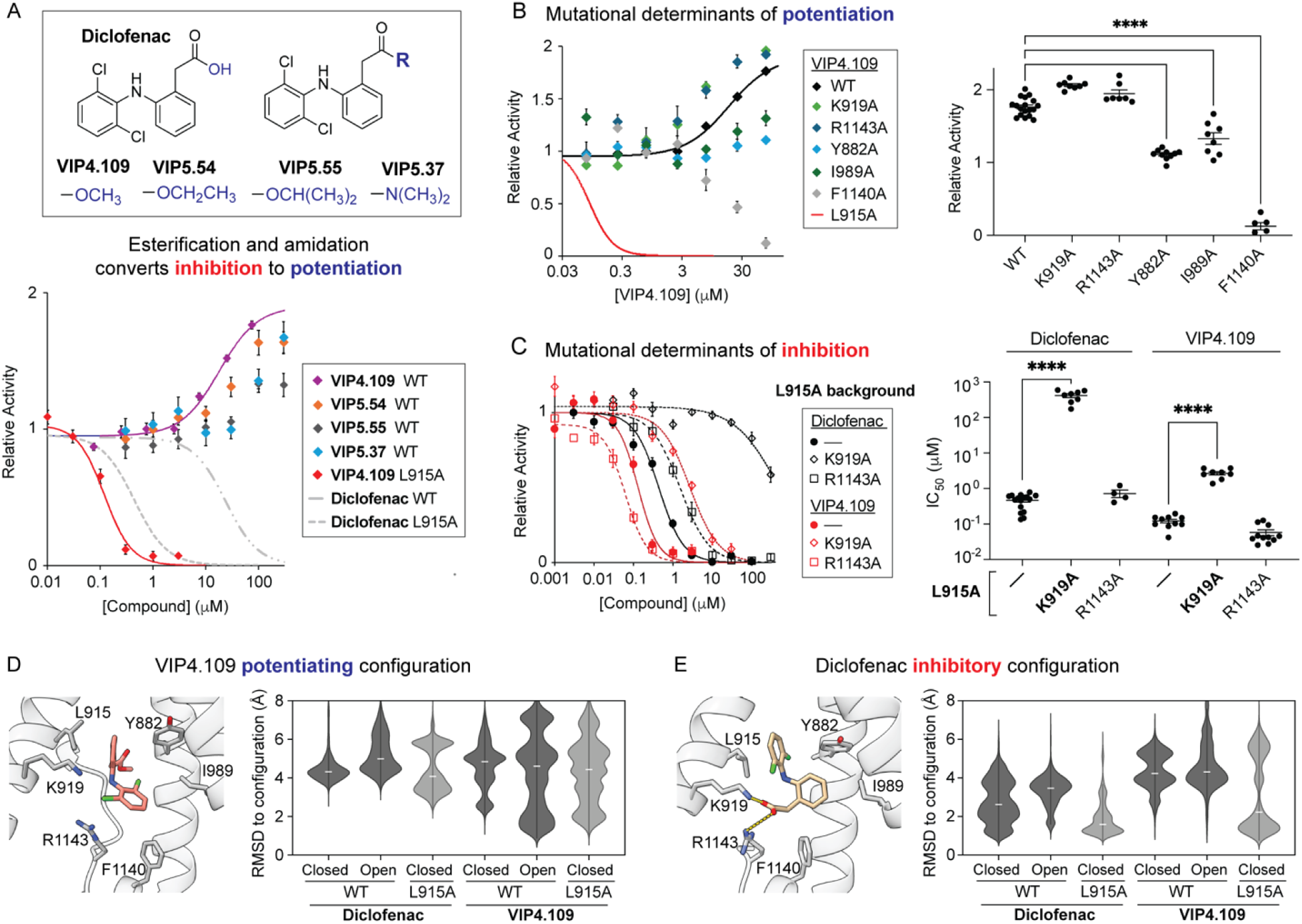
Modification of the diclofenac carboxylate switches efficacy between inhibition and potentiation. **(A)** Chemical structures of diclofenac ester and amide derivatives (top), and concentration–response relationships for modulation of PS-evoked TRPM3 activity by these compounds. TRPM3 activity was measured using a calcium mobilization assay in response to 50 μM PS. Data are shown as mean ± SEM (n=12-18). Dashed curves indicate inhibition of WT and L915A TRPM3 by diclofenac reproduced from Fig. 3 for comparison. **(B)** Concentration–response relationships for VIP4.109 modulation of WT and mutants. WT and K919A were activated by 50 μM PS, R1143A by 10 μM PS, Y882A and F1140A by 3 μM PS, and I989A by 1 μM PS. The summary plot shows relative activity measured at 75 μM VIP4.109. Data are shown as mean ± SEM (n=4-18). Statistical significance was determined by one-way ANOVA with Dunnett’s multiple-comparisons test relative to WT. **** p < 0.0001. **(C)** Concentration–response relationships for inhibition by diclofenac and VIP4.109 in the indicated mutant backgrounds. Summary plots show IC_50_ values as mean ± SEM (n=4-19). **(D)** Candidate potentiating configuration of VIP4.109 derived from MD simulations of the WT activated structure. Interacting residues are shown as sticks. Violin plots summarize the distribution of MD conformations relative to the candidate potentiating configuration. For clarity, only simulations initiated from configuration 2 are shown. **(E)** Candidate inhibitory configuration derived from MD simulations of diclofenac in the closed/inhibited TRPM3 structure. Violin plots summarize the distribution of MD conformations relative to the candidate inhibitory configuration. For clarity, only simulations initiated from configuration 1 are shown.

**Figure 5:**
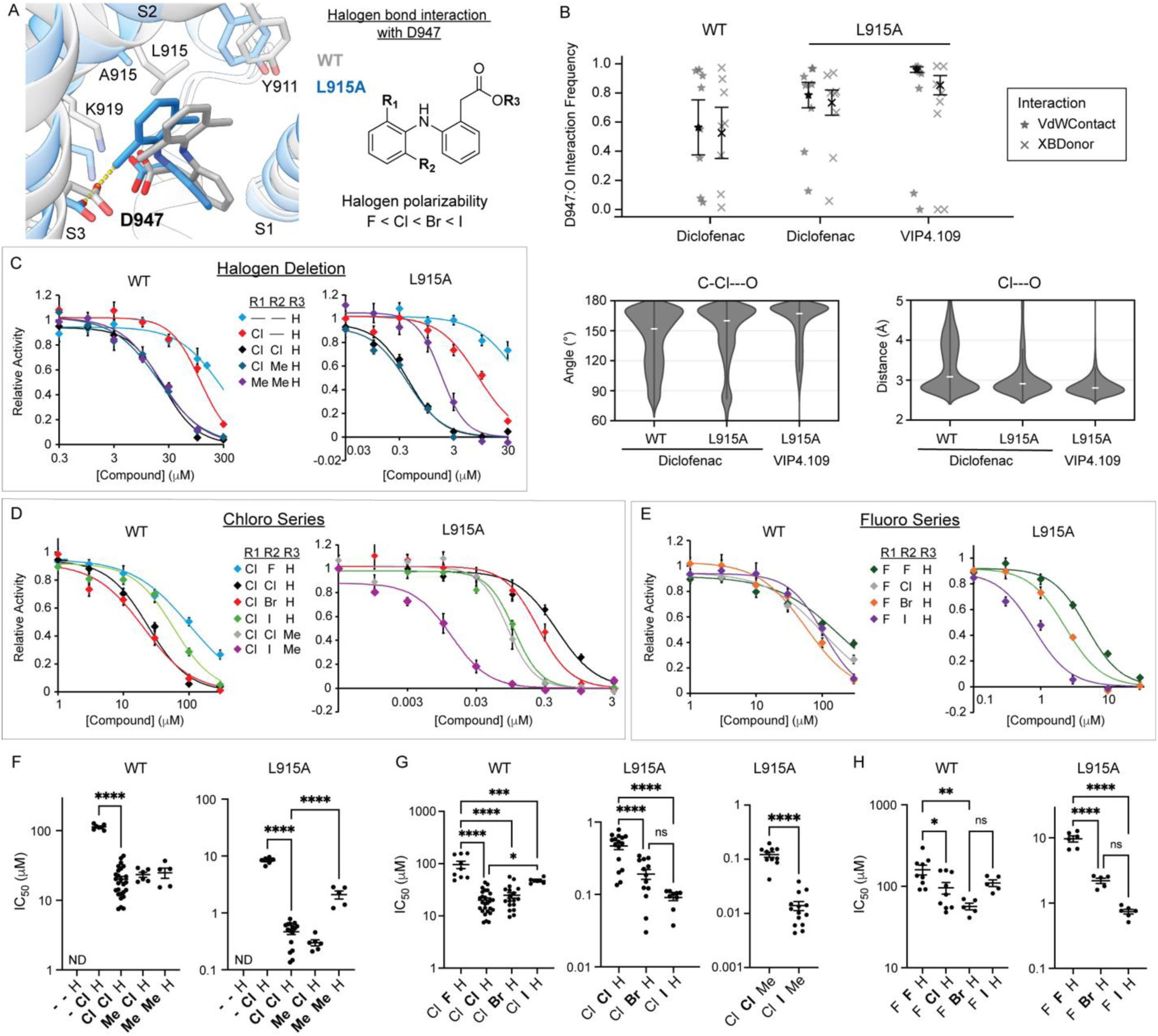
Structure–activity relationships support a halogen bond contributing to diclofenac potency. **(A)** The candidate inhibitory configuration of diclofenac in WT and L915A TRPM3 derived from molecular dynamics (MD) simulations. A chlorine is positioned near D947, consistent with a putative halogen bond. To the right is the chemical scaffold used for structure– activity analysis, with substitutions introduced at positions R1, R2 and R3. **(B)** Interaction frequencies between a chlorine atom in diclofenac or VIP4.109 and the oxygen in D947 from the indicated MD simulations initialized from configuration 1, considering all conformations within 3 Å of the candidate inhibitory pose. Shown are individual replicates with mean ± SEM. XBDonor indicates a halogen bond interaction. Below are violin plots showing the distribution of angles between C-Cl in the ligand and the oxygen in D947, and the interaction distance between Cl in the ligand and the oxygen in D947 for the indicated simulations. **(C–E)** Concentration–response relationships for inhibition of PS-evoked (50 μM) WT and L915A TRPM3 activity by diclofenac derivatives. Data are shown as mean ± SEM (n=5-29). The R1=F, R2=Cl, R3=H derivative unexpectedly potentiated L915A (Supplementary Fig. 17). **(F–H)** Summary of IC_50_ values for data in panels C–E shown as mean ± SEM. Statistical significance was determined by one-way ANOVA with Dunnett’s multiple-comparisons test. * p<0.05, ** p<0.01, *** p<0.001, **** p<0.0001.

### Diclofenac carboxylate derivatives switch efficacy between inhibition and potentiation

The MD simulations of diclofenac predict that the carboxylate forms ionic interactions with K919, R985, and R1143. To probe the dependency of this group, we tested the methyl ester derivative VIP4.109. Unexpectedly, VIP4.109 potentiated WT TRPM3 responses to PS (Fig. 4A). Ethyl ester (VIP5.54), propyl ester (VIP5.55), and dimethyl amide (VIP5.37) derivatives also potentiated WT TRPM3 (Fig. 4A), whereas the primary amide (VIP5.52) and monomethyl amide (VIP5.53) derivatives did not (Supplementary Fig. 12A). In contrast, VIP4.109 and the amide derivatives strongly inhibited L915A, with a potency rank order of primary amide (IC_50_ 10 nM) >> monomethyl amide ∼ methyl ester > dimethyl amide (Fig. 4A, Supplementary Fig. 12C). Among the ester derivatives, inhibitory activity decreased with increasing alkyl chain length: the ethyl ester was a weak inhibitor of L915A, whereas the propyl ester showed no detectable inhibition. Thus, modification of the diclofenac carboxylate can profoundly alter efficacy and potency, with neutral derivatives supporting either potentiation or strong inhibition depending on ligand chemistry and channel background.

Mutational analysis further indicated that inhibition and potentiation arise through distinct modes of VSLD engagement (Fig. 4B–C, Supplementary Fig. 13). Y882A and I989A reduced VIP4.109 potentiation, whereas F1140A converted potentiation to inhibition. In contrast, K919A strongly reduced inhibition by diclofenac and VIP4.109 in the L915A background but had little effect on VIP4.109 potentiation in the WT background. Thus, inhibition and potentiation exhibit overlapping but distinct residue dependencies within the same VSLD pocket.

To assess whether these differential functional effects are associated with distinct ligand configurations, we analyzed MD simulations of diclofenac and VIP4.109 initiated from both configurations in the cryo-EM structure (Supplementary Table 2). Simulations were performed using closed structures of WT and L915A and a previously reported putative activated WT structure (PDB 9B2A). The candidate inhibitory configuration identified in Fig. 3A was preferentially sampled in L915A (Fig. 4E). In the candidate inhibitory configuration of VIP4.109 in L915A, the methyl ester extends toward the solvent-exposed intracellular region of the VSLD pocket (Supplementary Fig. 12D), consistent with the ability of the amide derivatives to also support inhibition in L915A. In contrast, a distinct VIP4.109 configuration (i.e. a candidate potentiating configuration), characterized by a flipped orientation of the dichlorophenyl ring within the VSLD, was preferentially sampled in the activated WT structure (Fig. 4D, Supplementary Fig. 14 and 15). Together with the differential residue dependencies observed by mutagenesis, these findings support partially overlapping VSLD configurations associated with inhibition and potentiation.

### Structure-activity relationships support a halogen bond that contributes to diclofenac potency

The candidate inhibitory configuration revealed a putative halogen bond that may contribute to inhibitor potency. In this configuration, a diclofenac chlorine is positioned near D947, with D947 approaching along the C–Cl axis, a geometry consistent with a halogen bond (Fig. 5A and 5B). K919 and R985 also contact the chlorine but approach more laterally (Supplementary Fig. 16). The frequency of the D947-Cl interaction in the MD simulations increased from diclofenac/WT to diclofenac/L915A and was greatest for VIP4.109/L915A (Fig. 5B), paralleling the increased inhibitory potency of these ligand–channel combinations. The frequent sampling of this halogen bond in the MD simulations prompted us to systematically vary the halogen substituents in diclofenac to test whether halogen chemistry contributes to inhibitory potency.

Removal of both chloro substituents (-/-) or a single chloro substituent (Cl/-) significantly reduced inhibitory potency for both WT and the L915A mutant (Fig. 5C). In contrast, replacement of one chloro substituent with a methyl group (Cl/Me) preserved potency, and substitution of both chloro substituents with methyl groups (Me/Me) selectively reduced potency in L915A. These results suggest that the electronic properties of at least one chloro substituent contribute to inhibition in the L915A background where diclofenac preferentially occupies the inhibitory configuration.

Because halogen bond strength generally increases with halogen polarizability (F < Cl < Br < I), we next tested whether inhibitory potency followed this relationship. Increasing halogen polarizability systematically increased inhibitory potency in L915A in two independent analog series: Cl/Cl < Cl/Br < Cl/I and F/F < F/Br < F/I (Fig. 5D–H, Supplementary Fig. 17). This relationship was not retained in WT, consistent with the higher frequency and more favorable geometry of the halogen bond in L915A (Fig. 5B). The same principle extended to VIP4.109. Replacement of chlorine with iodine increased the inhibitory potency of VIP4.109 in L915A by ∼10-fold (Fig. 5D and 5G), consistent with the prediction that strengthening this interaction should enhance inhibition. Together, these results support a model in which halogen bonding contributes to inhibitory potency, particularly under conditions that favor the candidate inhibitory configuration.

## Discussion

Recent structures have established the TRPM3 VSLD as a major site of modulation by chemically diverse agonists and inhibitors^15–17^. Here, cryo-EM, MD simulations, mutagenesis, and medicinal chemistry define molecular determinants of maprotiline and diclofenac recognition and show how ligand-specific interactions within this adaptable allosteric pocket influence potency and efficacy.

Isosakuranetin, maprotiline, diclofenac, and primidone occupy the same VSLD cavity yet engage distinct subsets of residues. Rather than relying on a conserved pharmacophore, the VSLD therefore accommodates structurally diverse scaffolds through partially overlapping interaction networks (Fig. 6). This concept has important implications for ligand discovery efforts in TRPM3. A recent virtual screening campaign successfully identified novel TRPM3 inhibitors using structural information from the VSLD pocket^20^. The current study suggests that this pocket is particularly amenable to structure-based drug discovery because it contains multiple interaction hotspots capable of recognizing chemically diverse ligands. The recurring involvement of residues such as Y882 and I989 across multiple ligands highlights potential focal points for future screening and optimization efforts. The configurational heterogeneity observed in MD simulations further suggests that ensemble-based screening incorporating multiple receptor and ligand configurations may better capture the diversity of interactions supported by this pocket than screening against a single structure.

**Figure 6:**
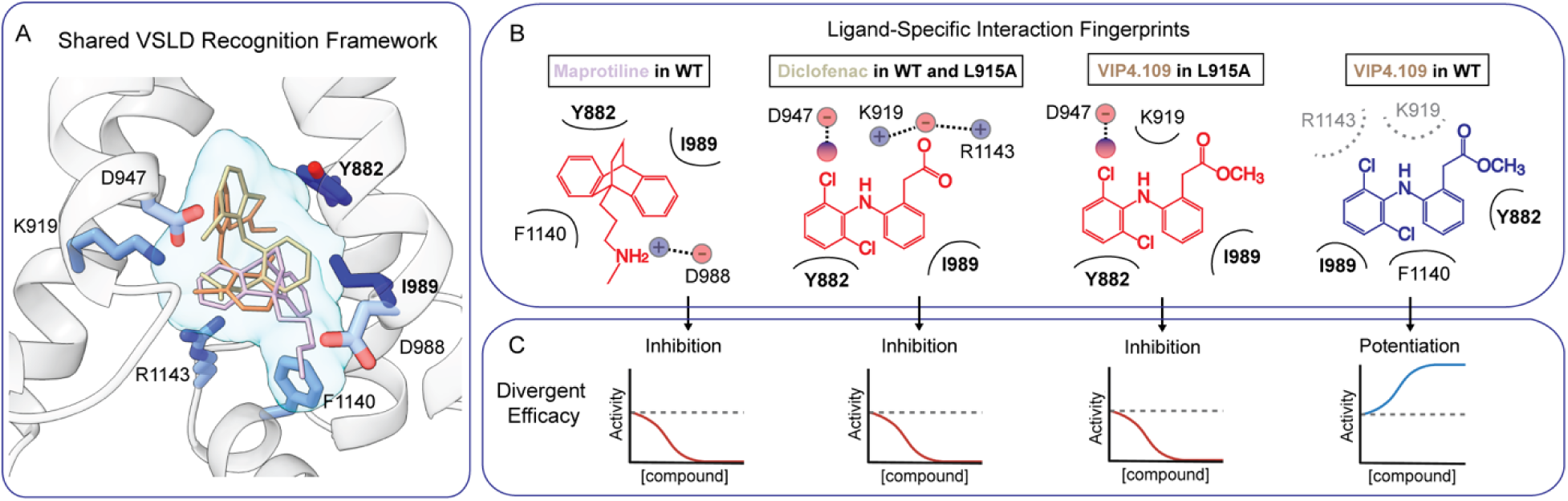
Framework for ligand recognition and efficacy in the TRPM3 VSLD. **(A)** Shared VSLD recognition framework. Superposition of representative VSLD-binding ligands (maprotiline, diclofenac, and VIP4.109) illustrates a common binding cavity within the TRPM3 voltage sensor-like domain. Residues identified through mutagenesis, molecular dynamics simulations, and structure–activity analyses as contributors to ligand modulation are shown as sticks. Despite their structural diversity, these ligands occupy overlapping regions of the VSLD, defining a shared recognition framework for TRPM3 modulators. **(B)** Ligand-specific interaction fingerprints. Simplified schematics summarize representative interactions underlying modulation by maprotiline, diclofenac, and VIP4.109 in WT and L915A TRPM3. I989 and Y882 are shared determinants that contribute to recognition of multiple ligand classes, whereas D988, D947, K919, R1143, and F1140 contribute in a ligand-or efficacy-dependent manner. **(C)** Divergent efficacy. Distinct interaction fingerprints within the VSLD are associated with different functional effects. Maprotiline and diclofenac inhibit WT TRPM3, whereas VIP4.109 potentiates channel activity. In L915A, VIP4.109 inhibits channel activity and exhibits altered residue dependence. Together, these observations support the model of an adaptable allosteric pocket, in which ligand-specific interactions and configurations shape efficacy.

Beyond defining ligand-specific interactions within the VSLD, our findings reveal that subtle changes in ligand chemistry can alter the direction of TRPM3 modulation. Notably, this efficacy switch is produced by modification of ligand chemistry alone, demonstrating that closely related compounds can produce opposing effects through the same VSLD pocket. VIP4.109 potentiation was reduced by Y882A and I989A, whereas F1140A converted potentiation to inhibition. In contrast, K919A strongly impaired inhibition but preserved VIP4.109 potentiation. These differential residue dependencies indicate that inhibition and potentiation involve partially overlapping modes of ligand engagement within the VSLD. Consistent with this interpretation, MD simulations identified distinct ligand configurations that were differentially sampled in closed and putative activated TRPM3 structures. Although the simulations do not establish a direct structural basis for the differential effects of individual mutations such as K919A, the combined functional and computational data support a model in which ligand efficacy is associated with redistribution among alternative configurations within the VSLD.

The diclofenac derivatives further identify halogen chemistry as an important determinant of inhibitory potency. Halogen bonds have become increasingly recognized as important determinants of ligand recognition in medicinal chemistry^21^, but have not been implicated experimentally in modulation of TRP channels. Aspartate and glutamate carboxylates can serve as halogen bond acceptors in protein-ligand complexes, with crystallographic and computational studies demonstrating favorable interactions between ligand chlorine atoms and aspartate carboxylate oxygens^22, 23^. MD simulations predict a putative halogen bond between diclofenac and D947. Although D947A was nonfunctional and therefore precluded direct mutational testing, the structure-activity relationships support this model: potency in L915A increased with halogen polarizability in two independent analog series, consistent with the expected energetic trend for halogen bonding. Steric and hydrophobic effects may also contribute, but the combined computational and chemical data support a measurable contribution of halogen bonding to ligand recognition within the VSLD.

In summary, the TRPM3 VSLD is an adaptable allosteric pocket that recognizes diverse ligands through distinct interaction fingerprints. Our results define molecular determinants of maprotiline and diclofenac inhibition, support halogen bonding as a contributor to inhibitor potency, and demonstrate that subtle chemical modifications can switch ligand efficacy. Functional and computational analyses further support redistribution among alternative VSLD configurations as a molecular basis for these opposing effects. Together, these findings provide a framework for understanding and structure-guided manipulation of TRPM3 pharmacology.

## Methods

### Expression and Purification of TRPM3

Mouse TRPM3α2 (residues 1-1,344) in a pEG BacMam vector was a generous gift from Professor Chen Zhao (University of Florida). The construct has a C-terminal PreScission protease cleavage site followed by an eGFP tag. Expression and purification of this protein was performed as previously reported with some modifications^14^. Briefly, DH10emBacVSV competent cells were transformed with the mouse TRPM3α2-eGFP plasmid to produce Bacmid DNA, which was used to transfect Sf9 cells^24^. Baculovirus production was monitored by mCherry signal from Sf9 cells, and P2 generation virus was harvested and used to transduce HEK293 GnTI^-^ cells at a density of 3 x 10^6^ with 10% baculovirus. 6 h after adding virus, sodium butyrate was added to 10 mM and the cells were maintained at 30°C for 48 h before harvesting.

Purification of mouse TRPM3α2 was performed using pellets from 4L of cells. The cells were resuspended in 150 ml of buffer (20 mM Na-Bicine pH 8.5, 500 mM NaCl, 150 mM KCl, 0.5 mM MgCl_2_, 2 mM CaCl_2_, and 2 mM Na_2_ATP) with protease inhibitor (AEBSF, aprotinin, leupeptin, pepstatin A, benzamidine, trypsin inhibitor) and DNAase (Worthington). DTT was omitted from the purification, in contrast to the original report^14^, in order to preserve a disulfide bond in the pore loop that is critical for TRPM3 structure and function^17, 19^. The resuspended cells were lysed by dounce homogenizer, centrifuged at 39,000xg for 30 min to recover membranes, and the membrane pellets homogenized in the same buffer. Extraction was performed using 1.2% GDN (Anatrace) for 1 h at 4°C, and insolubilized material removed by centrifugation at 39,000xg for 30 min. Affinity purification was performed by binding to in-house GFP-Nb-Sepharose resin for 1 h at 4°C, followed by two washes: 1) 30 column volumes of 20 mM Na-Bicine pH 8.5, 500 mM NaCl, 150 mM KCl, 100 mM MgCl_2_, 2 mM Na_2_ATP and 0.006% GDN, and 2) 10 column volumes of 20 mM Na-Bicine pH 8.5, 500 mM NaCl, 2.5 mM Na_2_ATP and 0.006% GDN. The protein was eluted by digestion with HRV3c protease for 1.5 h, and the eluate was further purified by size exclusion chromatography on a Superose 6 Increase column (10/300 GL, Cytiva) pre-equilibrated with 20 mM Na-Bicine pH 8.5, 500 mM NaCl and 0.006% GDN. Pooled fractions were concentrated to ∼3 mg/ml for cryo-EM.

### Cryo-EM sample preparation, imaging, and single particle analysis

The TRPM3 samples were prepared by adding the inhibitors 30 min prior to vitrification. The sample was kept on ice during this incubation. For each sample, 3 μl of sample was applied to a freshly plasma-cleaned Quantifoil R1.2/1.3 300 mesh gold grid. Grids were treated with a hydrogen/oxygen plasma for 60 seconds using a Gatan Solaris 950 system immediately prior to sample preparation. Following sample deposition, grids were blotted for 2 seconds at 95% relative humidity at 18°C before being plunge-frozen in liquid ethane with a Vitrobot Mark IV (Thermo Fisher Scientific). Images were collected on a Krio 300 kV Cryo-EM equipped with a Falcon 4 Direct Electron Detector. Single particle cryo-EM data was acquired using counting mode on the Falcon 4 with the EPU software 3.8.1. For all datasets, movies were collected with a pixel size of 0.776 Å. For the isosakuranetin and maprotiline datasets, movies were collected with a total dose of 55 electrons per Å^2^, and for the diclofenac dataset, a total dose of 51 electrons per Å^2^.

Single particle cryo-EM datasets were processed in CryoSPARC v4.7.0^25^ and RELION-5^26^ (Supplementary Figure 1). The general workflow started with cryoSPARC using patch motion and CTF correction, particle selection using blob picker followed by Topaz picking^27^, and multiple rounds of 2D classification. Ab initio volumes served as references for multiple rounds of heterogeneous refinement, and the final reconstructions in cryoSPARC were produced using non-uniform refinement. These particles were then further processed in RELION-5 using 3D classification. For the isosakuranetin and maprotiline dataset, 3D classification focused on the TMD was performed without alignment, while for the diclofenac dataset, 3D classification of the whole protein was performed with alignment. The optimal workflow was chosen based on the approach that yielded the best local resolution in the VSLD.

For all three TRPM3 structures, PDB 8ED7 was used as an initial model, and real space refinement was performed in PHENIX 1.19.2^28^. The structures were manually built into the cryo-EM density map using COOT 0.9.6^29^, refined in PHENIX and this process was iteratively repeated. Ligands and phospholipids were retrieved from the PDB database, and geometry or cif files were optimized using the grade web server (https://grade.globalphasing.org). 3D volume visualization and figure preparation was performed using ChimeraX 1.10.

### Calcium mobilization assay

TRPM3 function was assessed using a calcium mobilization assay. Full-length mouse TRPM3α2 with an N-terminal GFP, along with all mutations, were purchased from Genscript (Accession: NM_001366141.2). The calcium mobilization assay was performed using a 96-well plate format in a Flexstation3 (Molecular Devices). Human embryonic kidney (HEK293T) cells were maintained in minimum essential medium (MEM) media supplemented with L-glutamine, 100 U/ml penicillin and streptomycin and 10% fetal bovine serum. HEK293T cells were transfected with the full-length mouse TRPM3α2-GFP WT or mutant DNA (1,600 ng) and GCaMP6 (6.4 μg) using Lipofectamine 2000 (Invitrogen). After 24 h, the transfected cells were seeded at 50,000 cells in a volume of 100 μL in poly-D-lysine coated black-wall clear-bottom 96-well plates. After 48 h, the MEM media was exchanged for Hank’s Balanced Salt Solution (ThermoFisher Cat No. 14025076) three times. Calcium mobilization or influx into the cells was measured at 25°C in the Flexstation-3 using an excitation wavelength of 485 nm and fluorescence emission detection at 525 nm. The standard protocol involved measurement of baseline fluorescence signal for 30 s, followed by injection of pregnenolone sulfate (PS) at 50 μM and measurement of TRPM3 response as indicated by an increase of fluorescence (i.e. calcium influx) for 130 s. Subsequently, 2 μM ionomycin was injected to determine the maximum response. To measure the inhibitory or potentiating effects of compounds, each compound was added to each well at the desired concentration ∼5 min prior to reading the plate. Then with each injection of PS, the test compound was also included to keep the concentration of that compound constant. In all cases, PS and inhibitors were prepared in DMSO stocks and the final concentration of DMSO added to the each well in the plate was kept below 0.1%. For each condition, 2-4 replicates were performed within a single plate, and this was repeated for two or more transfections. The normalized activity (reported as normalized fluorescence-F) for each condition was determined by subtracting the background fluorescence signal (samples in which only HBSS was injected) from the PS-induced response, and dividing this by the ionomycin response. The normalized activity/fluorescence in the presence of inhibitor at different concentrations was divided by the normalized activity in the absence of inhibitor yielding the relative activity, and this was plotted against ligand concentration to produce concentration response curves. These were fit to the following function to calculate an IC_50_ for half-maximal inhibition:

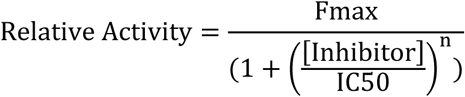

### Molecular dynamics simulations

All-atom simulations of ligand-bound TRPM3 were carried out for five different assemblies: maprotiline-bound WT, diclofenac-bound WT, diclofenac-bound L915A, VIP4.109-bound WT and VIP4.109-bound L915A. The maprotiline-bound cryo-EM structure was used as an initial model for the maprotiline simulations, while the diclofenac-bound cryo-EM structure was used for the diclofenac- and VIP4.109-bound simulations. The starting configurations of VIP4.109 ligands were modeled to align with the diclofenac configurations. For the diclofenac-bound WT and VIP4.109 WT assemblies we also used the previously solved cryo-EM structures of TRPM3 in a putative active conformation (PDB 9B2A)^17^, removing all previous ligands and adding diclofenac/VIP4.109 by aligning in turn each binding pocket of the active structure with the diclofenac-bound inactive structure. Systems were mutated in silico using ChimeraX^30^. We used the entire cryo-EM structures of the maprotiline-bound and the diclofenac-bound WT and L915A systems in the inactive state. To balance computational efficiency with physiological relevance, we truncated the remaining systems at the N-termini positions 1-516 and at the C-termini positions 1161 to the end, as has been done previously^17^. Missing loops were modelled using MODELLER^31^, using the original protein sequence except for the larger gaps at positions 588-630 and 795-860 where glycine linkers of length 5 and 10, respectively, were used instead. Titratable residues were protonated to match pH 7.4, and the systems were then embedded in symmetric 1-palmitoyl-2-oleoyl-sn-glycero-3-phosphocholine (POPC) membrane bilayers and subsequently solvated in TIP3P water and 150 mM NaCl using the membrane builder in CHARMM-GUI^32–34^. The systems were minimized and equilibrated using the standard procedure in CHARMM-GUI, with gradual release of positional restraints on protein, membrane and ligands. All simulations were performed using GROMACS 2024^35^, utilizing the CHARMM36 forcefield^36^ to describe the protein with ligand parameters generated using CGenFF in CHARMM-GUI^32, 37^. The temperature was kept at 300 K using a velocity-rescaling thermostat^38^, and the pressure was kept constant at 1 bar using a stochastic cell rescaling barostat^39^. The particle mesh Ewald algorithm was used for long-range electrostatic interactions, and hydrogen-bond lengths were constrained using the LINCS algorithm. A force-based smoothing function was employed for pairwise nonbonded interactions at 1 nm with a cutoff of 1.2 nm. Bonded and short-range nonbonded interactions were calculated every 2 fs, and periodic boundary conditions were employed in all three dimensions. Production runs consisted of four replicates for the maprotiline-bound system and two sets of two replicates for each diclofenac- and VIP4.109-bound systems, with each set corresponding to both ligand orientations consistent with the experimental density. All replicates were run independently for 500 ns, and frames were analyzed every 100 ps.

All four ligands per replicate were analyzed independently, resulting in 16 ligand-protein systems for the maprotiline-bound systems and 8 ligand-bound systems per ligand orientation and system for the diclofenac- and VIP4.109-bound systems. The ligands were aligned using the Cα atoms of the corresponding VSLD, delineated by the residues 870-892, 903-960, 971-994 and 1126-1144. Protein-ligand interactions were determined using ProLIF^40^ while water interactions and ligand positions, aligned relative to the VSLD, were determined using MDAnalysis^41^. The aligned positions were clustered over the 3× *N_atoms_*-dimensional space of the cartesian coordinates corresponding to the *N_atoms_*number of ligand atoms, across the simulation frames, using KMeans in Scikit-learn. The clusterings were visualized using principal component analysis.

## Data Availability

The data supporting the findings of this study are available within the paper and supplementary information files. The cryo-EM maps have been deposited in the Electron Microscopy Data Bank (EMDB) under accession codes EMD-76844, EMD-76843 and EMD-76429. The structural coordinates have been deposited in the RCSB Protein Data Bank (PDB) under the accession codes 12XW, 12XV and 12GV. MD simulation parameter files, trajectories, and representative poses have been deposited on Zenodo (https://doi.org/10.5281/zenodo.22955947).

## Acknowledgements

This study was supported by the International Anesthesia Research Society Frontiers in Anesthesia Research Award and NIH grant R35GM137957 to W.W.C., a post-doctoral AHA fellowship (24POST1189869) and Owens Anesthesiology Research Fellowship to V.D., and start-up funds from Washington University for S.M. Work by S.E.L., E.L. and R.J.H. was funded by grants from the Swedish Research Council (VR: 2019-02433, 2021-05806), Knut and Alice Wallenberg Foundation (KAW: 2023.0254), BioExcel-3 Centre-of-Excellence (EuroHPC Joint Undertaking: 101093290), Sven and Lilly Lawski’s Fund and Swedish eScience Research Center, with computational resources provided by the National Academic Infrastructure of Supercomputing in Sweden (NAISS: 2025/3-71). We are grateful to Dr. Chen Zhao for providing the construct and guidance for expression and purification of TRPM3.

## Authors Contributions

W.W.C. conceived and designed the project. V.D. led the experimental work and performed protein production, cryo-EM, and functional analyses. V.A.R. and S.M. designed and synthesized the diclofenac derivatives. S.E.L., R.J.H., and E.L. designed, performed, and analyzed the MD simulations. K.T.M. optimized protein expression and purification. All authors reviewed the manuscript.

## Competing Interests Statement

S.M. is a cofounder of Sparian biosciences.

