## Supplementary Information for "Molecular determinants of ligand recognition and efficacy in the TRPM3 voltage sensor-like domain"

Supplementary Figure 1

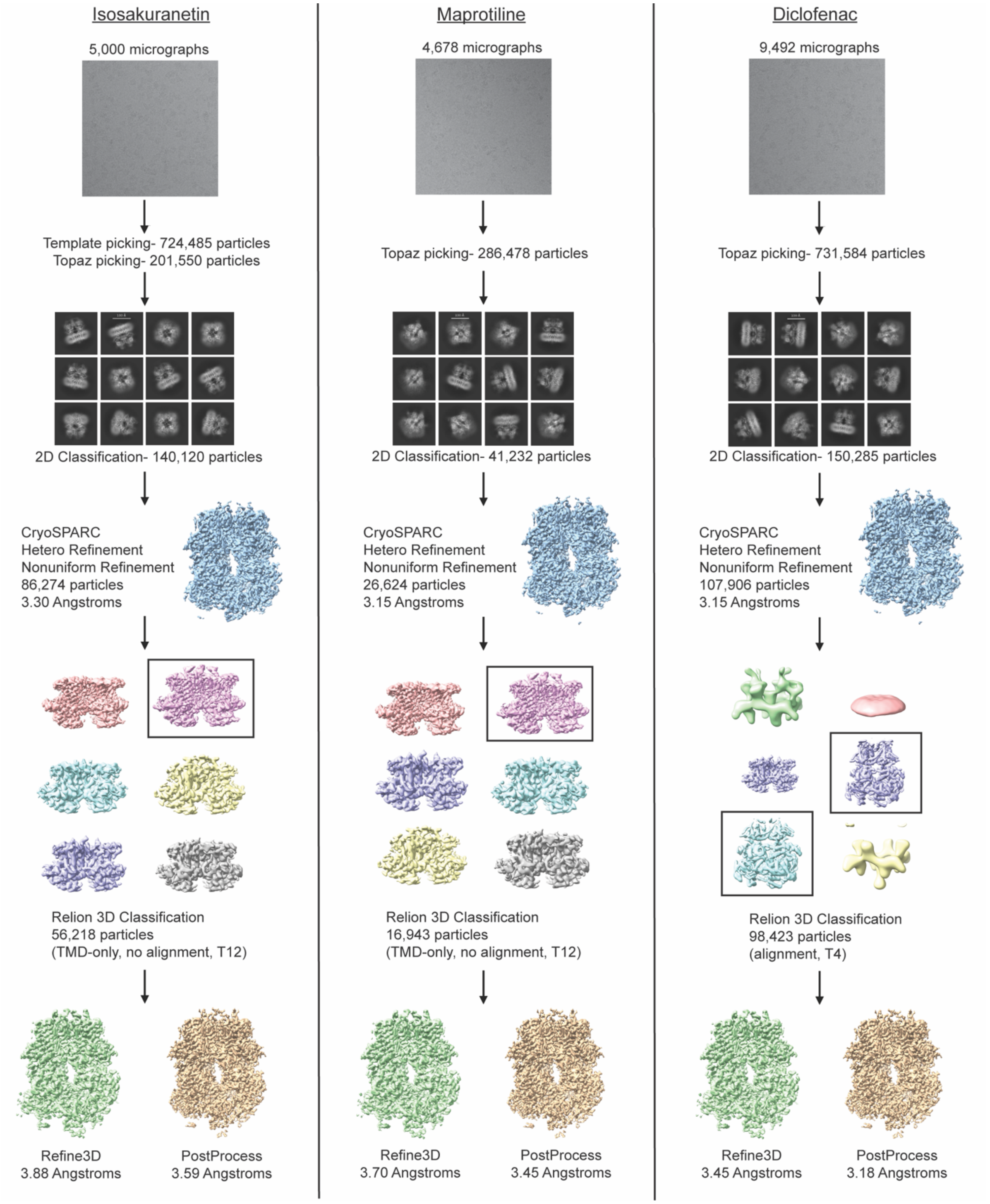


Supplementary Figure 1: Workflow of single particle analysis for TRPM3 structures showing the number of starting micrographs and the particles processed at each step. For the 3D classification jobs in Relion5, the box indicates the selected class, TMD-only refers to focused 3D classification with a mask encompassing the TMD, and the T value refers to the regularization parameter.

Supplementary Figure 2

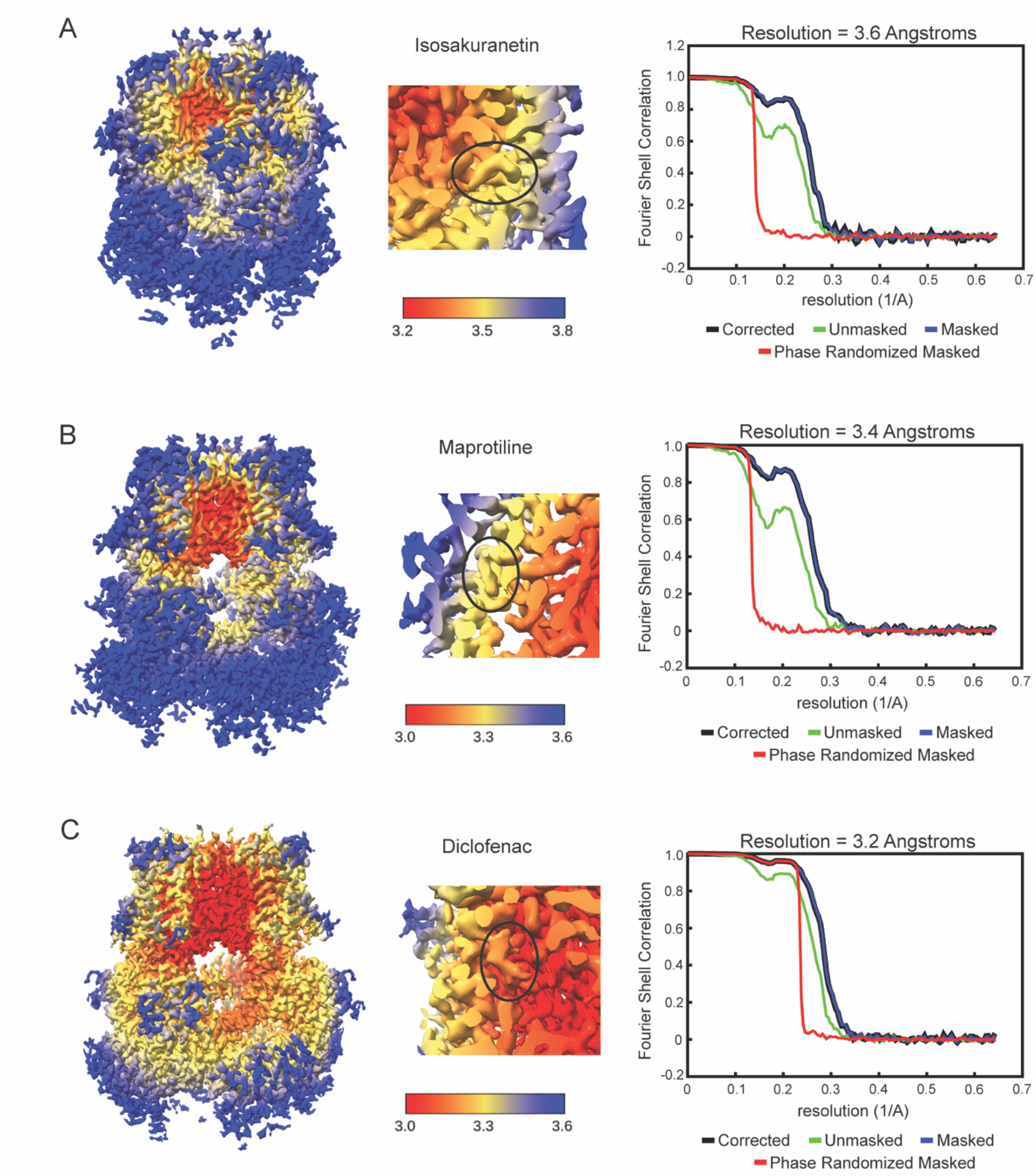


Supplementary Figure 2: The local and global resolution estimates for each structure shown as color representation on the cryo-EM density map, and Fourier Shell Correlation curves (FSC) from Relion5.

Supplementary Figure 3

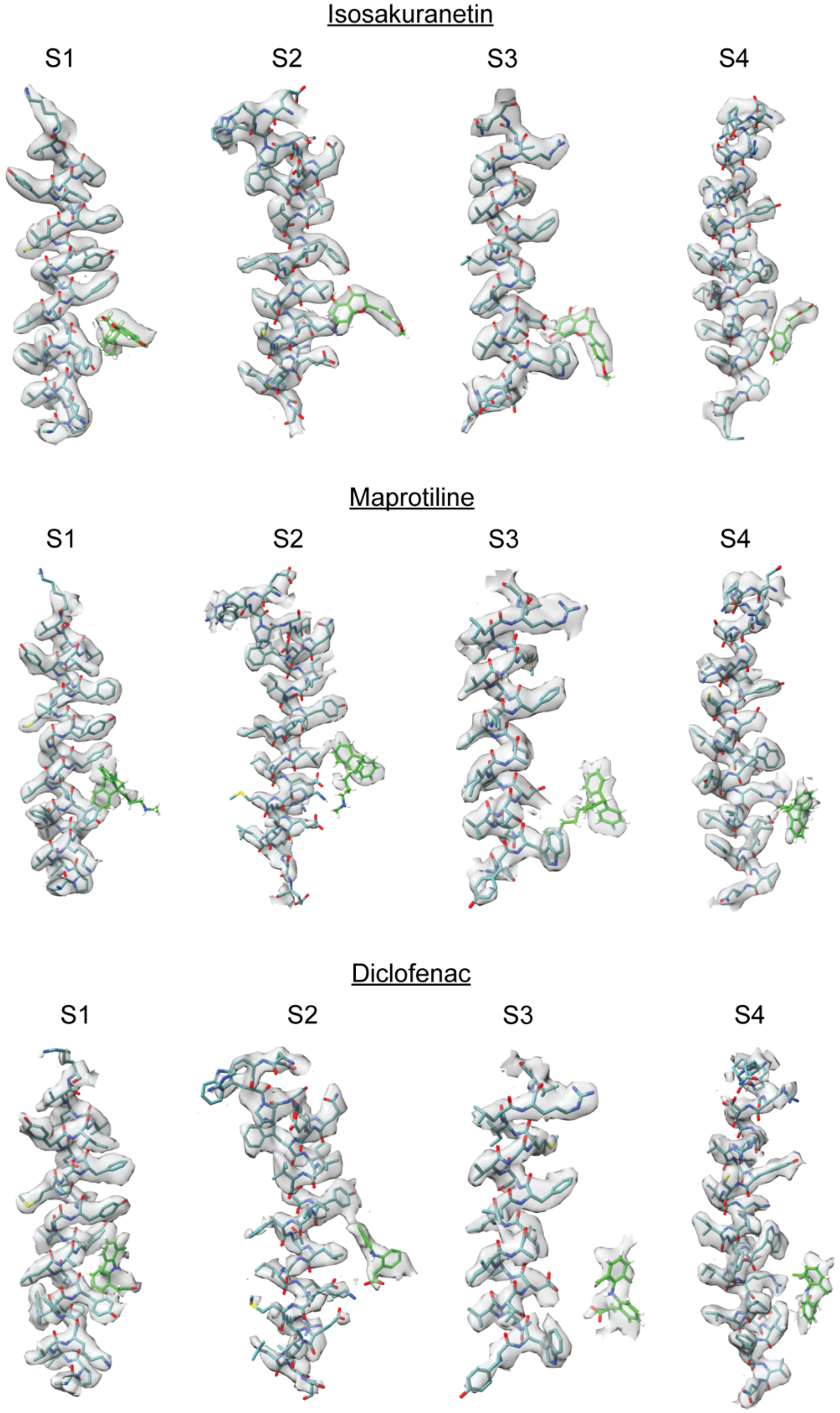


Supplementary Figure 3: Map-model fit for the S1-S4 helices in the VSLD and the ligand corresponding to isosakuranetin, maprotiline and diclofenac. The cryo-EM densities are set a sigma level of 3.0.

Supplementary Figure 4

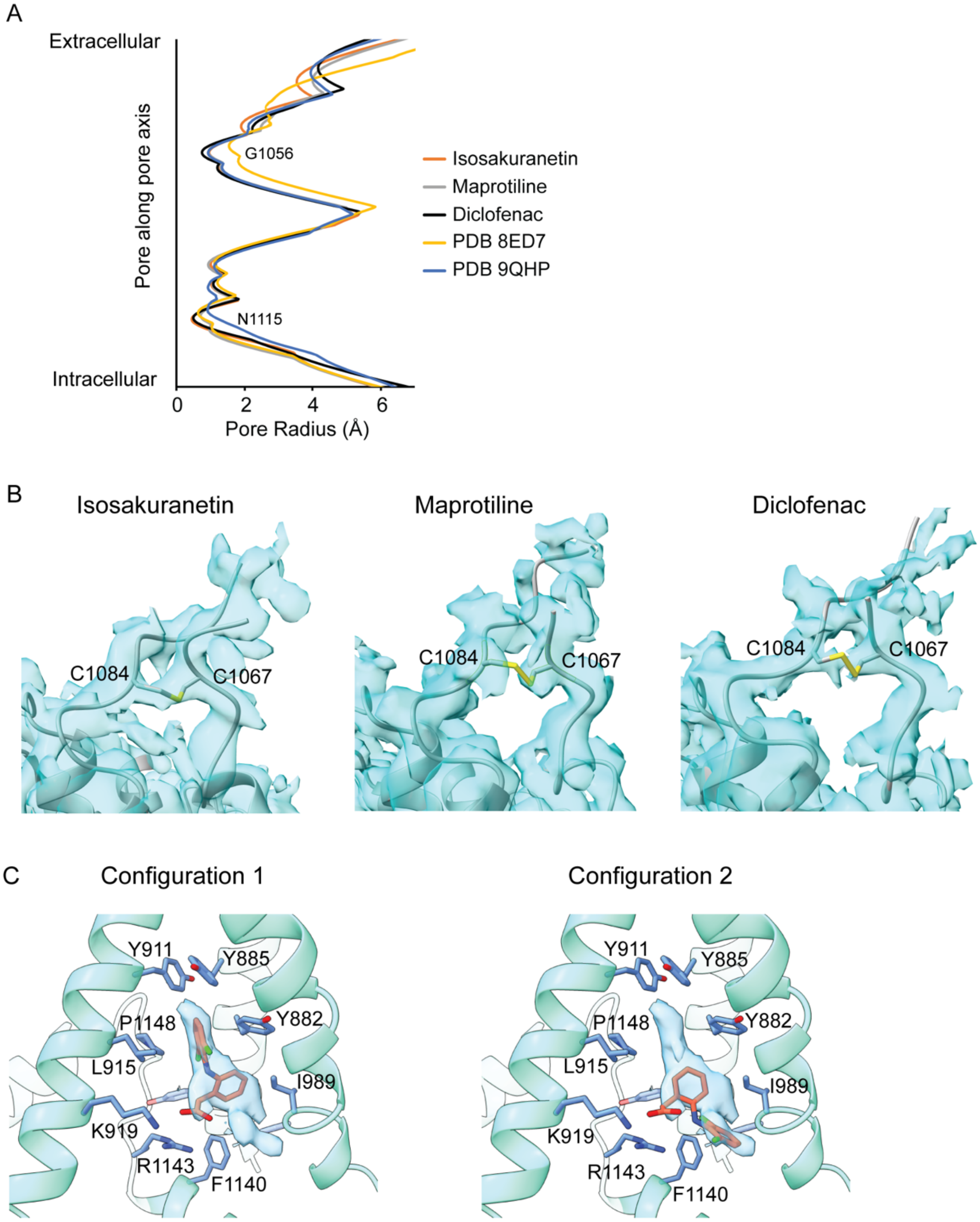


Supplementary Figure 4: (A) Plots of pore radius for the cryo-EM structures determined in the present study, and compared with structures of mouse TRPM3 in the apo condition (PDB 8ED7) and bound to isosakuranetin (PDB 9QHP). (B) Density and models for the isosakuranetin, maprotiline and diclofenac cryo-EM structures determined in the present study showing the extracellular pore loop that contains a disulfide bond (C1067 and C1084), most apparent in the isosakuranetin structure. (C) Cryo-EM density and two distinct proposed configurations for diclofenac within the VSLD cavity. Diclofenac is shown as sticks with the corresponding cryo-EM density. Side chains of residues within 4 Å of diclofenac are shown as sticks.

Supplementary Figure 5

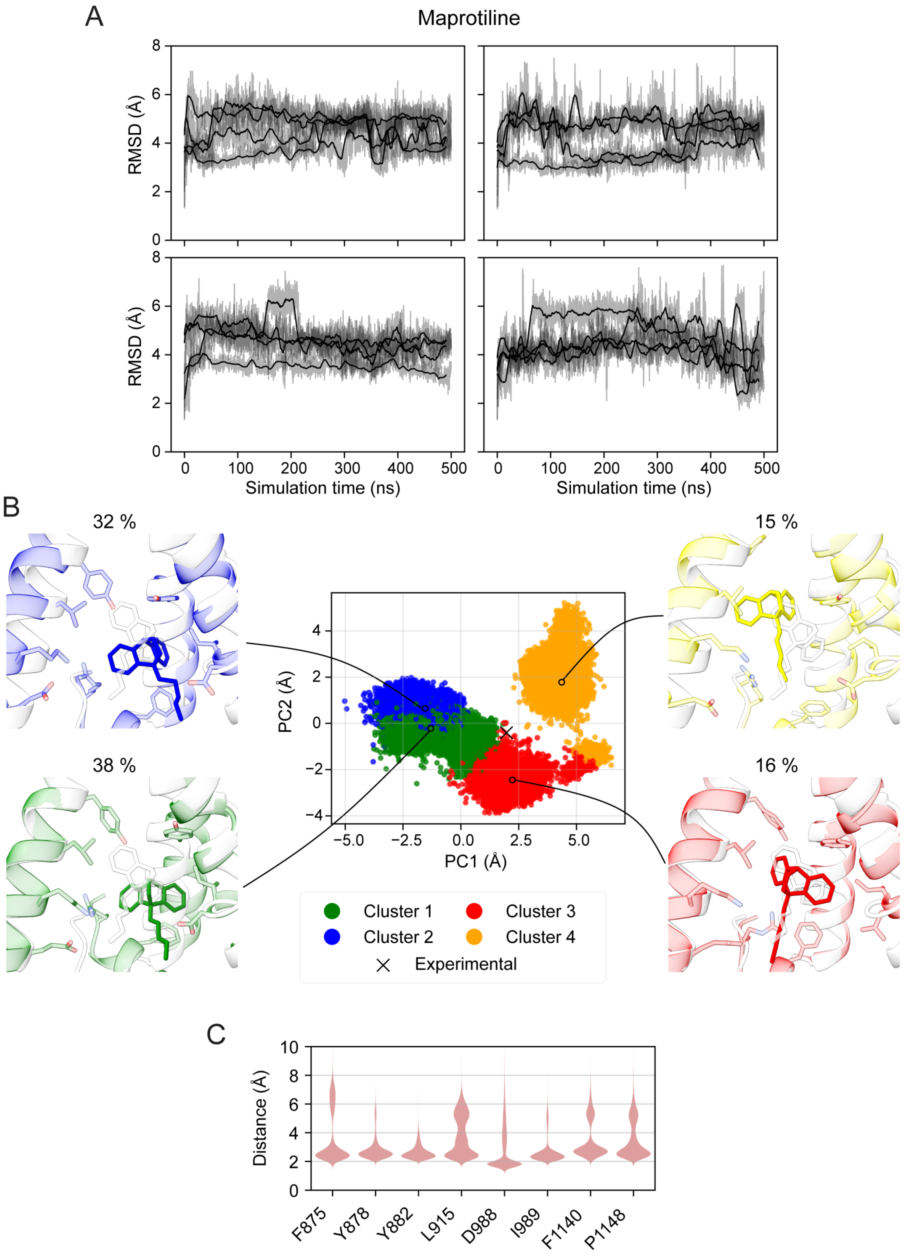


Supplementary Figure 5: Analysis of MD simulations of the TRPM3 structure with maprotiline. (A) RMSD of maprotiline compared to the experimental reference structure, aligned on the pocket represented by the Cα atoms of the surrounding VSLD helices. Each box corresponds to an independent replicate and each line to a single ligand. (B) Clustering of ligand positions when aligned on the VSLD. The center plot shows a projection of the aligned positions on the first two principal components, with colors indicating cluster assignments. Black circles correspond to the cluster centroids, and the corresponding ligand positions shown on the sides, with cluster population shown on top. (C) Violin plots highlighting the distribution of the minimum distance of maprotiline from select residues, across all maprotiline simulations.

Supplementary Figure 6

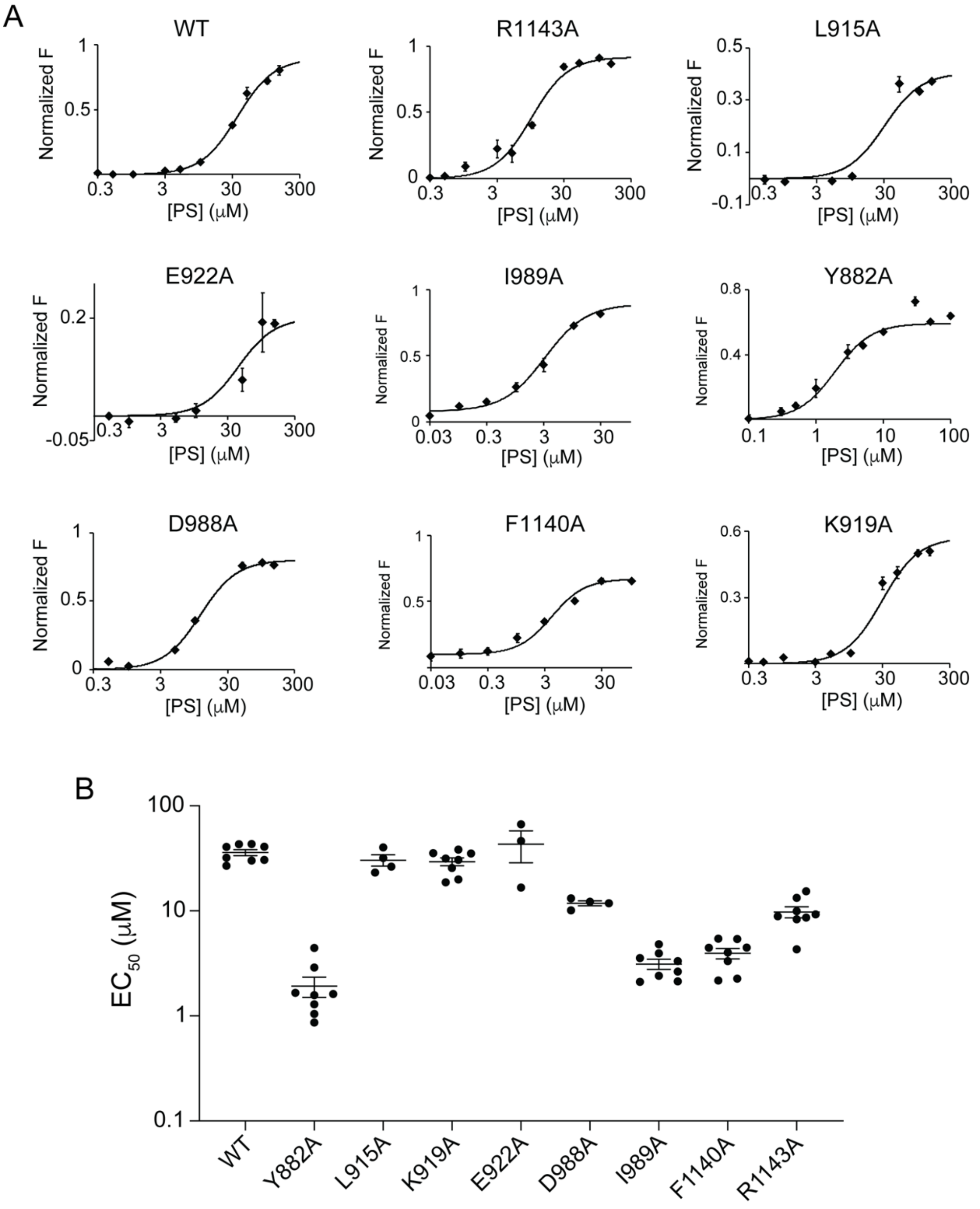


Supplementary Figure 6: Pregnenolone sulfate sensitivity of WT and mutant TRPM3 channels. (A) Concentration-response relationships for activation of WT and the indicated TRPM3 mutants by pregnenolone sulfate (PS), measured using a calcium mobilization assay. Responses were normalized to ionomycin. Data are shown as mean ± SEM. (B) Plot of the $EC_{50}$ for WT and each mutant derived from the data in panel (A), showing mean ± SEM.

Supplementary Figure 7

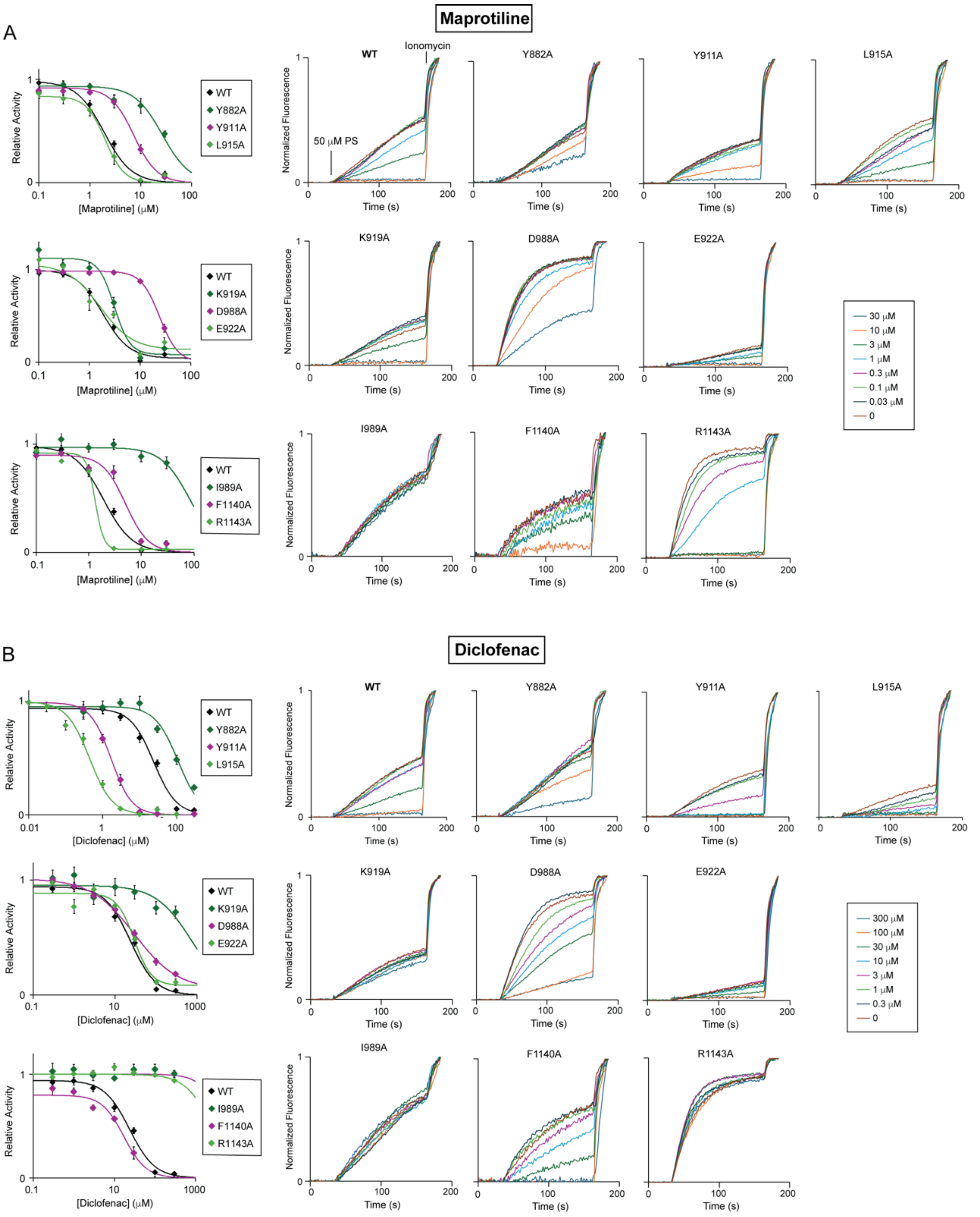


Supplementary Figure 7: Calcium mobilization data of WT and mutant TRPM3. The left shows concentration response curves for maprotiline and diclofenac inhibition of WT and mutants (n=4-

25, ±SEM). The right shows representative traces for each condition. The increase in fluorescence signal occurs in response to the injection of PS, and this is assayed in the presence of varying concentrations of maprotiline and diclofenac. Maximum response is determined by the addition of ionomycin, and each trace is normalized to the ionomycin response. 3 μM PS was used for I989A and Y882A, 5 μM PS for F1140A, and 50 μM PS for WT and the remaining mutants.

Supplementary Figure 8

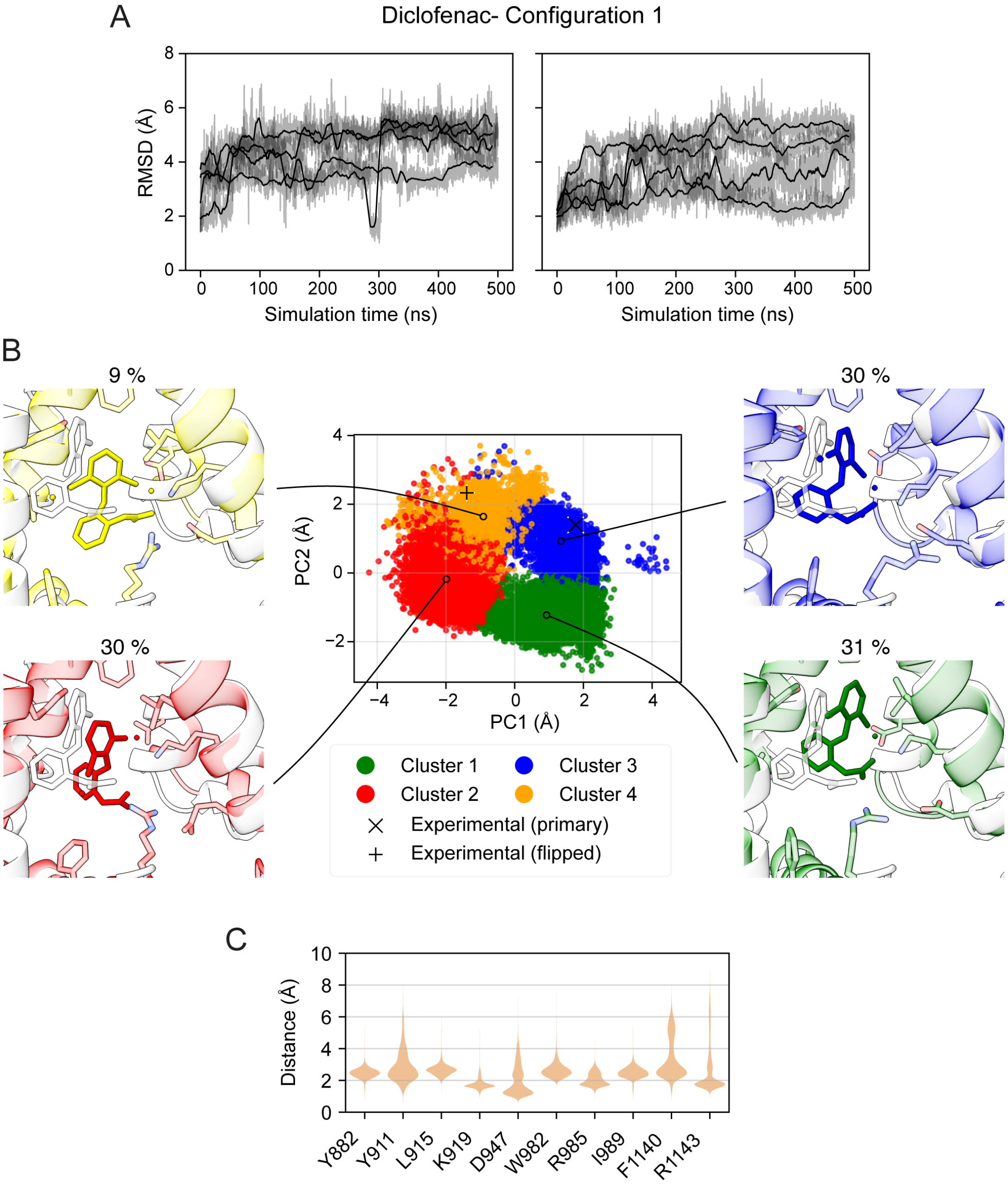


Supplementary Figure 8: Analysis of MD simulations of the TRPM3 structure with diclofenac starting in configuration 1. (A) RMSD of diclofenac compared to the corresponding experimental reference structure, aligned on the pocket represented by the Cα atoms of the surrounding VSLD helices. Each box corresponds to an independent replicate and each line to a single ligand. (B) Clustering of ligand positions when aligned on the VSLD. The center plot shows a projection of the aligned positions on the first two principal components, with colors indicating cluster assignments. Black circles correspond to the cluster centroids, and the corresponding ligand positions shown on the sides, with cluster population shown on top. (C) Violin plots highlighting the distribution of the minimum distance of diclofenac from select residues, across the WT diclofenac simulations initialized from configuration 1.

Supplementary Figure 9

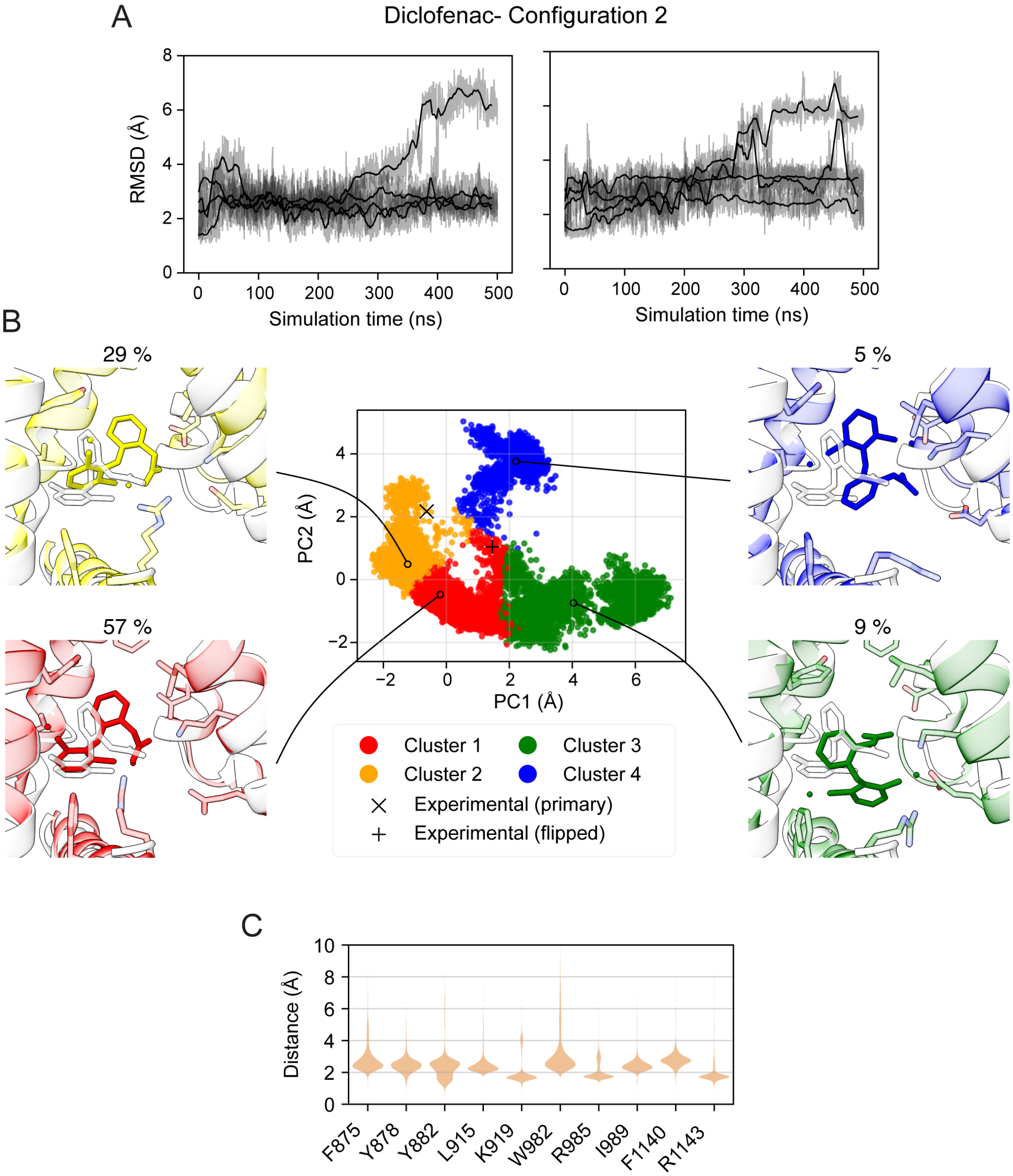


Supplementary Figure 9: Analysis of MD simulations of the TRPM3 structure with diclofenac starting in configuration 2. (A) RMSD of diclofenac compared to the corresponding experimental reference structure, aligned on the pocket represented by the Cα atoms of the surrounding VSLD helices. Each box corresponds to an independent replicate and each line to a single ligand. (B) Clustering of ligand positions when aligned on the VSLD. The center plot shows a projection of the aligned positions on the first two principal components, with colors indicating cluster assignments. Black circles correspond to the cluster centroids, and the corresponding ligand positions shown on the sides, with cluster population shown on top. (C) Violin plots highlighting the distribution of the minimum distance of diclofenac from select residues, across the WT diclofenac simulations initialized from configuration 2.

Supplementary Figure 10

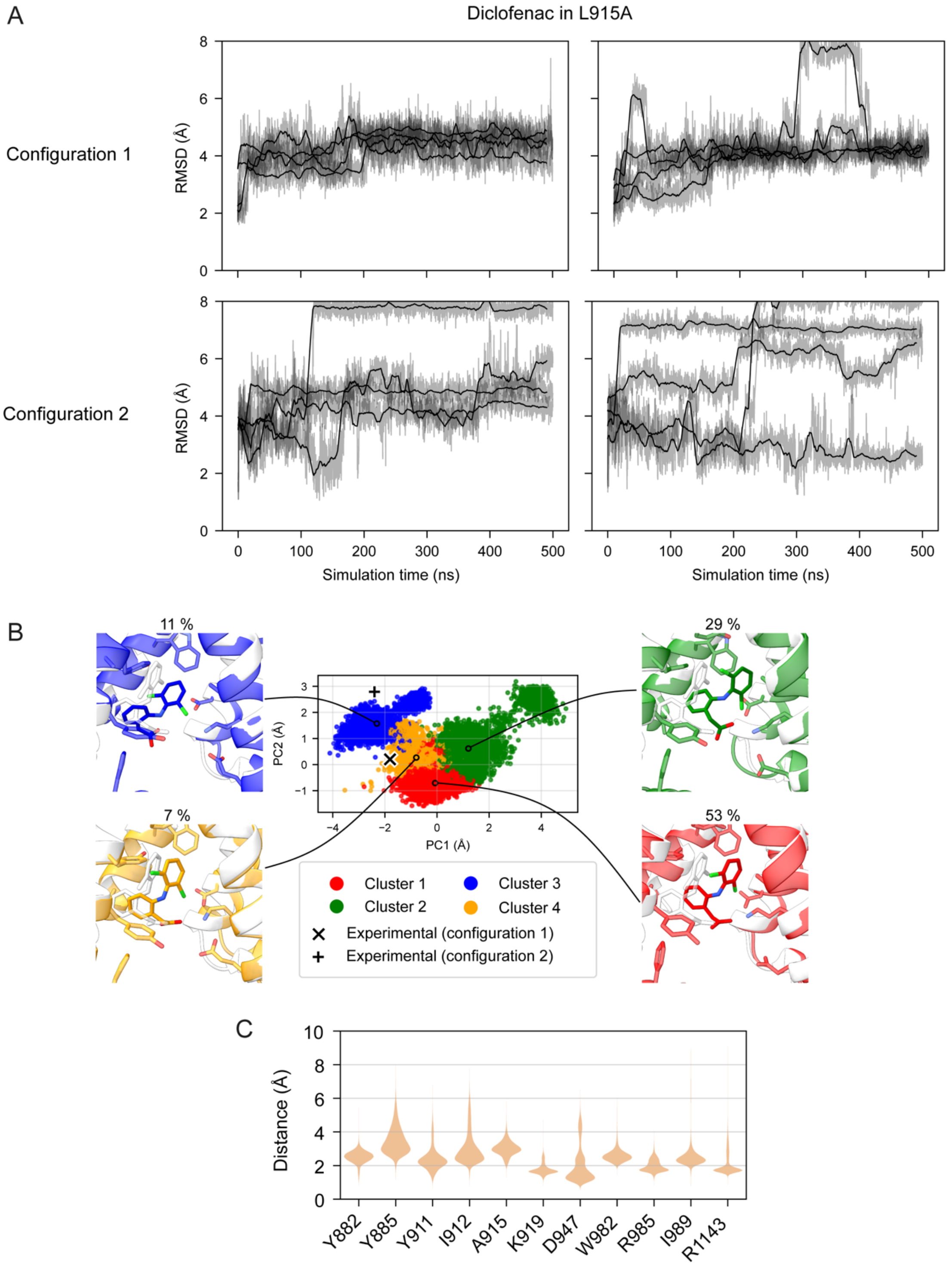


Supplementary Figure 10: Analysis of MD simulations of the L915A TRPM3 structure with diclofenac starting in configurations 1 and 2. (A) RMSD of diclofenac compared to the corresponding experimental reference structure, aligned on the pocket represented by the Cα atoms of the surrounding VSLD helices. Each box corresponds to an independent replicate initiated from configuration 1 (top) or configuration 2 (bottom), and each line corresponds to a single ligand. (B) Clustering of ligand positions when aligned on the VSLD, showing the results for the simulations starting in configuration 1. The center plot shows a projection of the aligned positions on the first two principal components, with colors indicating cluster assignments. Black circles correspond to the cluster centroids, and the corresponding ligand positions shown on the sides, with cluster population shown on top. (C) Violin plots highlighting the distribution of the minimum distance of diclofenac from select residues, across the L915A diclofenac simulations initialized from configuration 1.

Supplementary Figure 11

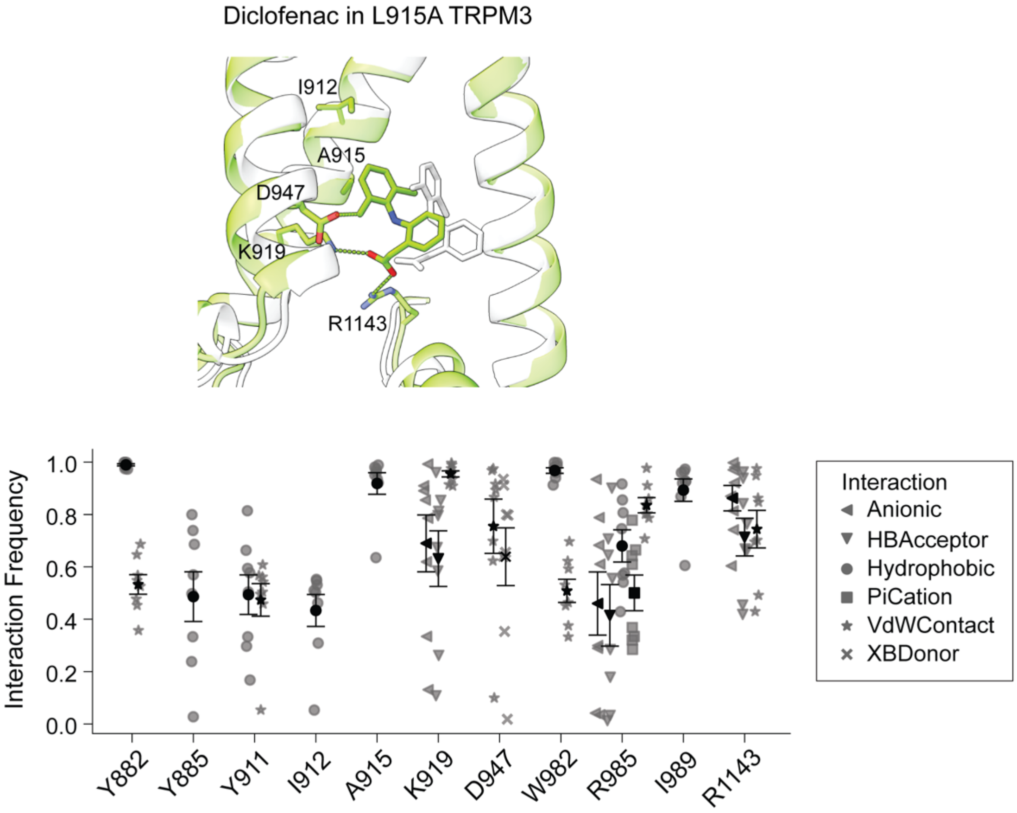


Supplementary Fig. 11: Representative pose of diclofenac in the L915A mutant simulations. Interaction frequencies between diclofenac and VSLD residues from the MD simulations of the L915A mutant initialized from configuration 1. Shown are individual replicates with mean ± SEM, showing all interactions with mean ≥0.4.

Supplementary Figure 12

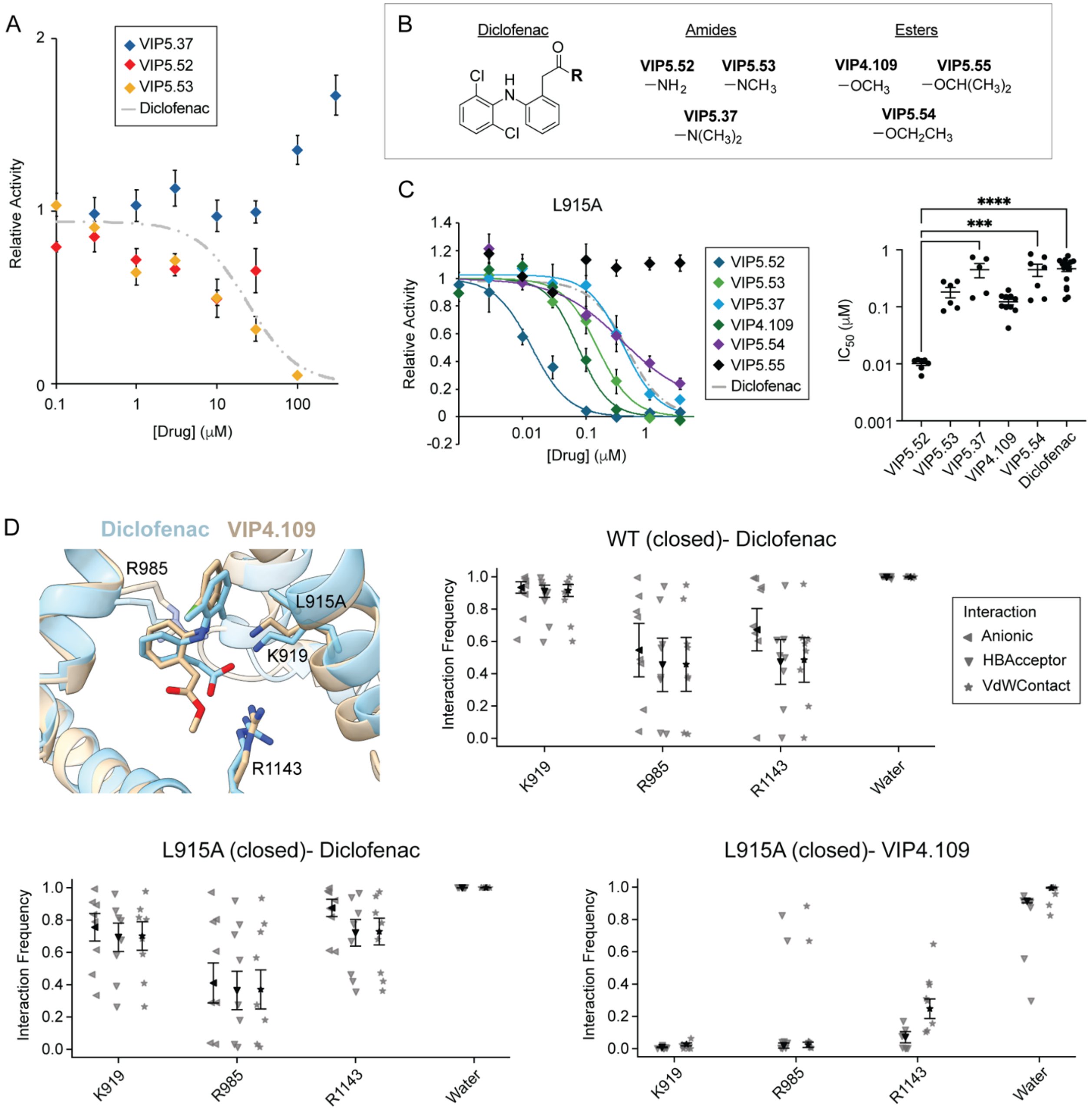


Supplementary Figure 12: Inhibition of L915A TRPM3 by ester and amide derivatives. **(A)** Concentration–response relationships for modulation of WT TRPM3 by diclofenac amide derivatives in response to 50 µM PS. Data are shown as mean ± SEM (n=12-16). **(B)** Chemical structures of diclofenac and ester or amide derivatives. **(C)** Concentration–response relationships for inhibition of L915A TRPM3 by diclofenac amide and ester derivatives in response to 50 µM PS, and plot of $IC_{50}$ values. Data are shown as mean ± SEM (n=5-10). Statistical significance was determined by one-way ANOVA with Dunnett's multiple-comparisons test. *** $p < 0.001$, **** $p < 0.0001$. (D) The candidate inhibitory configuration for diclofenac and VIP4.109 from MD simulations in the L915A closed structure, and plots of interaction frequencies for the indicated MD simulation systems initialized from configuration 1, considering all conformations within 3 Å of the candidate inhibitory pose. Shown are individual replicates with mean ± SEM.

Supplementary Figure 13

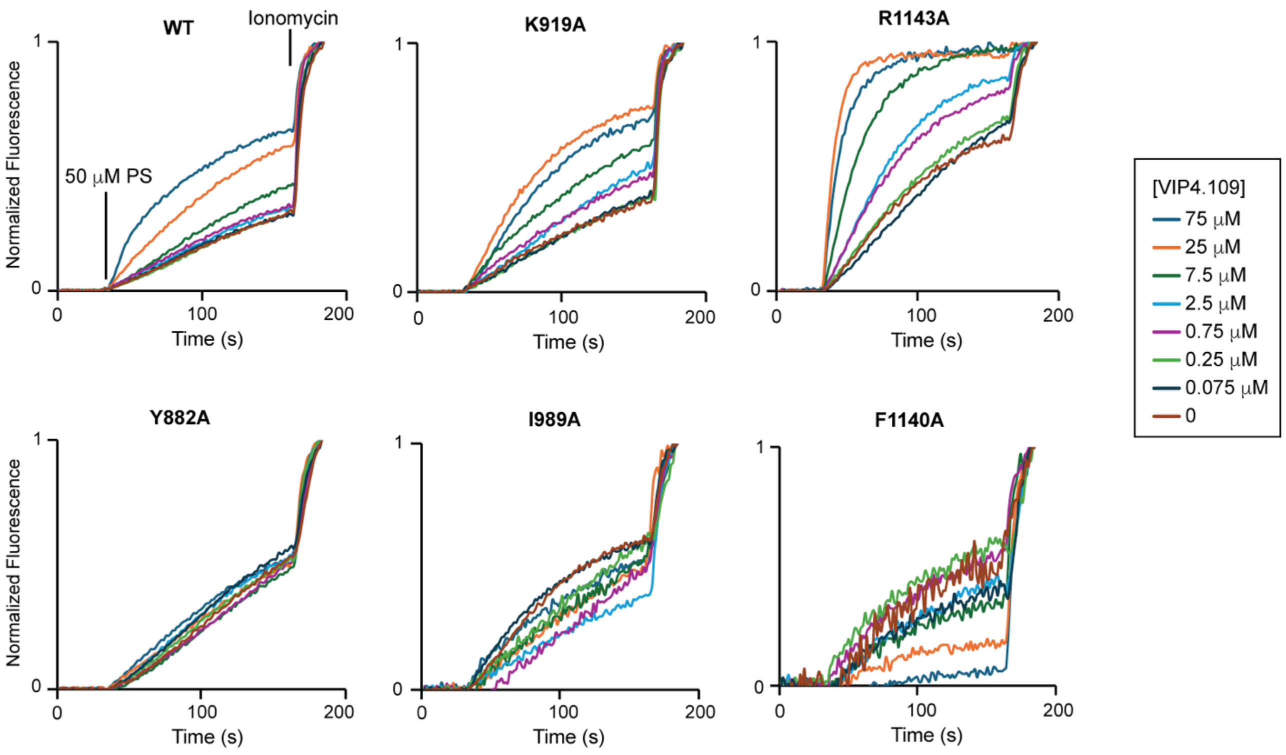


Supplementary Figure 13: Representative traces from calcium mobilization data of WT and mutant TRPM3 in response to PS and modulated by VIP4.109. The increase in fluorescence signal occurs in response to the injection of PS, and this is assayed in the presence of varying concentrations of VIP4.109. Maximum response is determined by the addition of ionomycin, and each trace is normalized to the ionomycin response. PS concentrations were chosen to achieve ~50% response relative to ionomycin to optimize evaluation of VIP4.109 potentiation and inhibition. 1 μM PS was used for I989A and Y882A, 3 μM PS for F1140A, 10 μM PS for R1143A, and 50 μM PS for WT and the remaining mutants.

Supplementary Figure 14

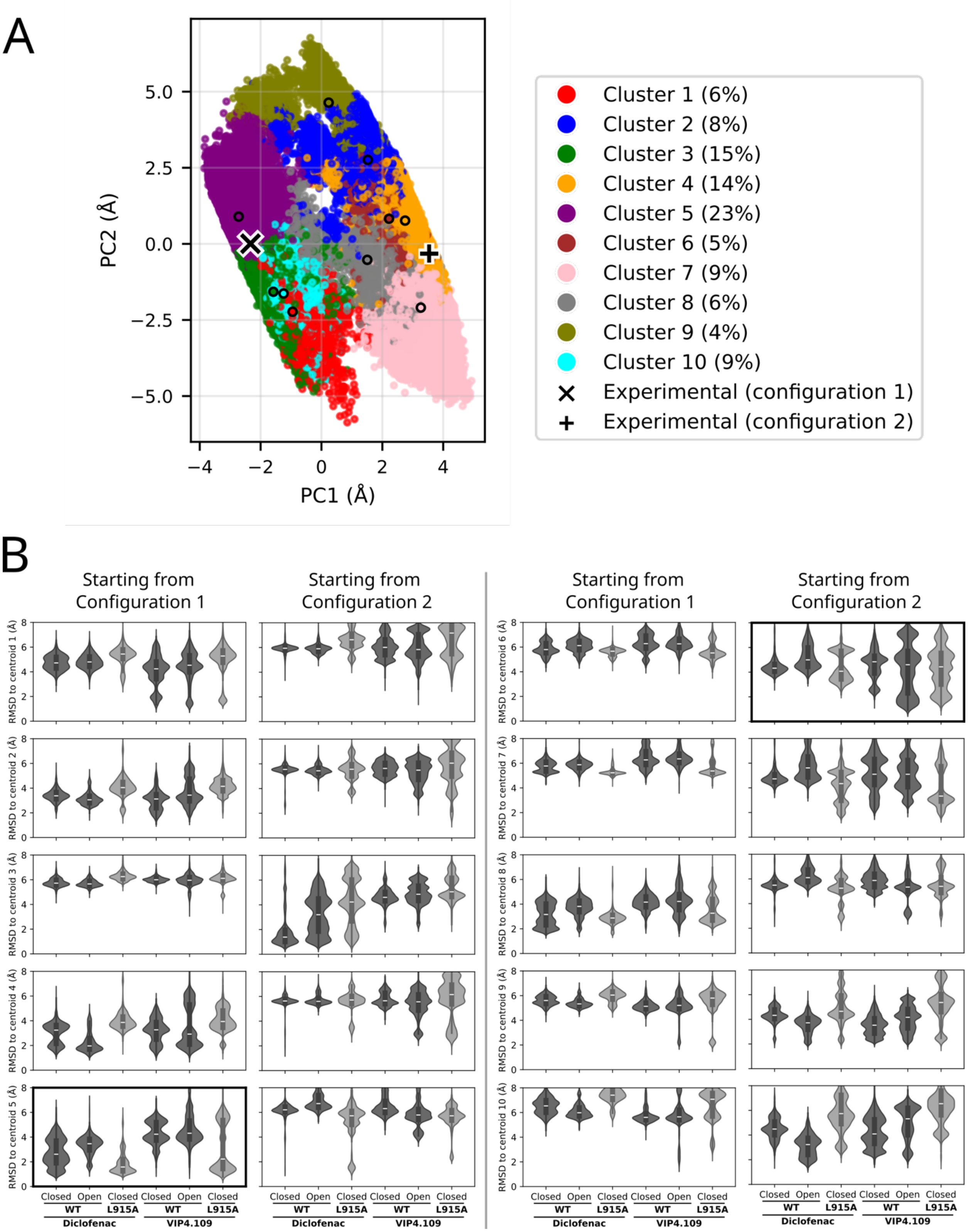

Supplementary Figure 14: Analysis of aggregate MD simulations of the WT and L915A TRPM3 structures (closed and open/activated for WT) with diclofenac and VIP4.109 starting in configurations 1 and 2. (A) Clustering of ligand positions when aligned on the VSLD. The left plot shows a projection of the aligned positions on the first two principal components, with colors indicating cluster assignments. Black circles correspond to the cluster centroids, and the experimental ligand positions and cluster populations are shown on the sides. Cluster 5 was chosen as the candidate inhibitory pose while cluster 8 was chosen as the candidate potentiating pose. (B) Violin plots summarize the distribution of MD conformations relative to the centroid for each cluster of 1-10 for simulations starting from configuration 1 and 2.

Supplementary Figure 15

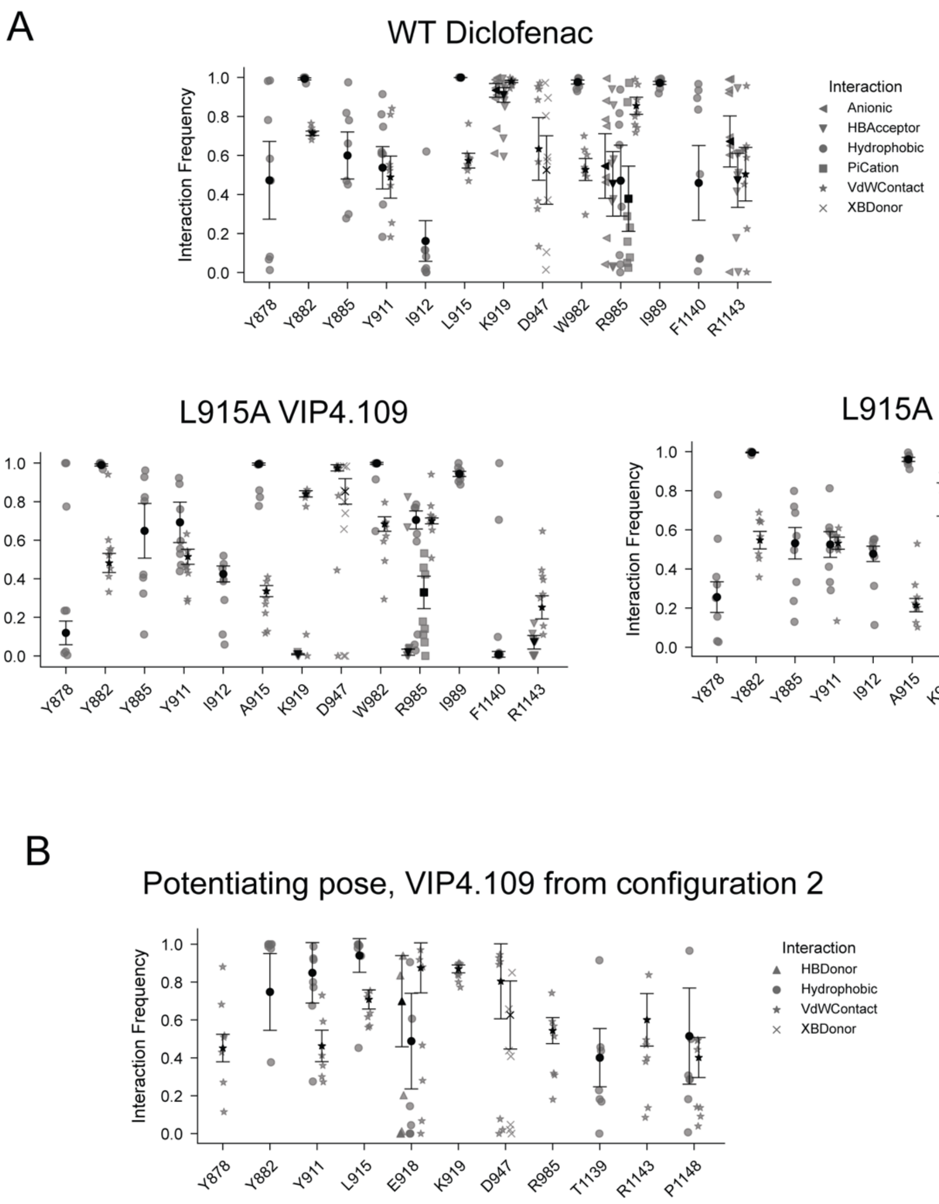


Supplementary Figure 15: (A) Interaction frequencies for conformations within 3 Å of the candidate inhibitory pose from simulations initiated from configuration 1. Shown are individual replicates with mean ± SEM, showing all interactions with mean ≥0.4. (B) Interaction frequencies for conformations within 3 Å of the candidate potentiating pose from the simulations initiated from configuration 2.

Supplementary Figure 16

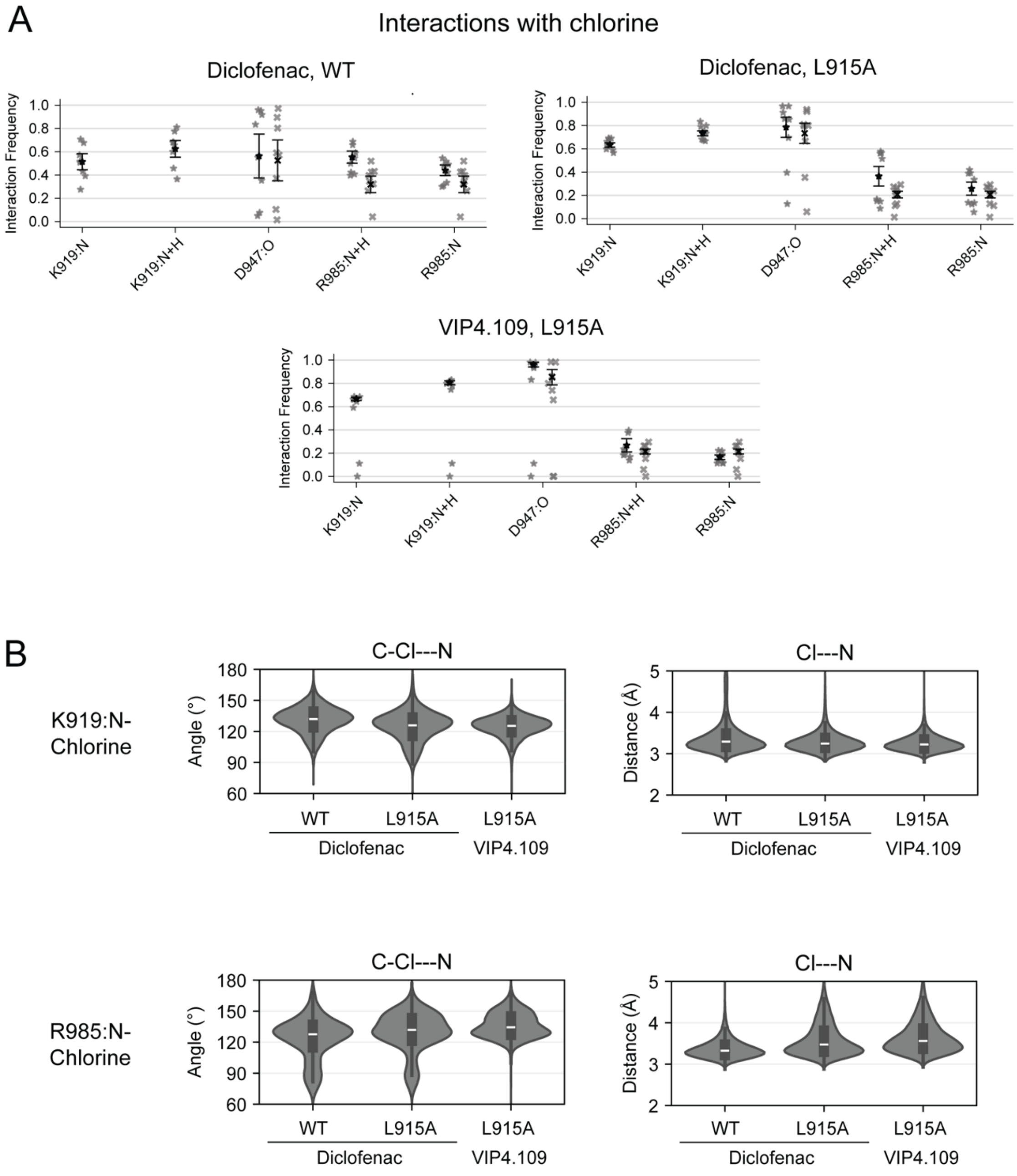


Supplementary Figure 16: (A) Interaction frequencies of K919, D947 and R985 with a chlorine in diclofenac/VIP4.109 for conformations within 3 Å of the candidate inhibitory pose from the indicated simulations. The data are from MD simulations initiated from configuration 1. Shown are individual replicates with mean ± SEM. (B) Violin plots showing the distribution of angles between C-Cl in the ligand and the nitrogens in K919 and R985, and the interaction distance between Cl in the ligand and the nitrogens in in K919 and R985 for the indicated simulations.

Supplementary Figure 17

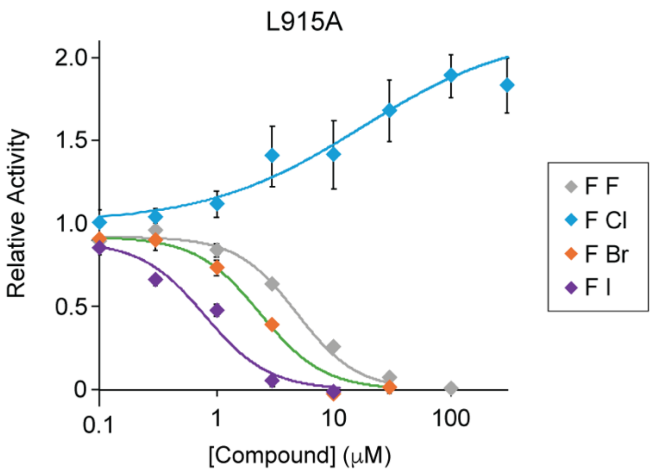


Supplementary Figure 17: Concentration response curve for the halogen series of diclofenac derivatives consisting of F/F, F/Cl, F/Br and F/I. The F/Cl derivative potentiates PS-evoked (50 μM) responses in the L915A TRPM3 mutant. Data are shown as mean ± SEM (n=8-12).

Supplementary Table 1: Cryo-EM data collection, refinement, and validation statistics

| | TRPM3 with Maprotiline EMD-76844 PDB 12XW | TRPM3 with Diclofenac EMD-76843 PDB 12XV | TRPM3 with Isosakuranetin EMD- 76429 PDB 12GV |
|---|---|---|---|
| Magnification | 75000 | 75000 | 75000 |
| Voltage (kV) | 300 | 300 | 300 |
| Electron exposure (e-/ Å$^2$) | 55.04 | 51 | 55.04 |
| Defocus range (μm) | 1.0 to 2.4 | 1.0 to 2.4 | 1.0 to 2.4 |
| Pixel size (Å) | 0.776 | 0.776 | 0.776 |
| Symmetry imposed | C4 | C4 | C4 |
| Initial particle images (no) | 286,478 | 731,584 | 926,035 |
| Final particle images (no) | 16,943 | 98,423 | 54,872 |
| Map resolution (Å) | 3.45 | 3.18 | 3.6 |
| FSC threshold | 0.143 | 0.143 | 0.143 |
| Initial model used | PDB 8ED7 | PDB 8ED7 | PDB 8ED7 |
| Model resolution (Å) | 3.4 | 3.2 | 3.7 |
| FSC threshold | 0.5 | 0.5 | 0.5 |
| Map sharpening B factor (Å$^2$) | -77.962 | -106.875 | -94.0569 |
| Non-hydrogen atoms | 31804 | 31796 | 31840 |
| Protein Residues | 3876 | 3876 | 3880 |
| Ligands | 12 | 12 | 12 |
| Protein | 30.45 | 38.67 | 22.68 |
| Ligand | 20.68 | 13.35 | 15.07 |
| Bond lengths (Å) | 0.006 | 0.005 | 0.004 |
| Bond angles (°) | 0.971 | 0.945 | 0.896 |
| MolProbity Score | 2.23 | 2.30 | 2.16 |
| Clashscore | 26.25 | 28.04 | 28.80 |
| Poor rotamers (%) | 0 | 0 | 0 |
| Favored (%) | 95.32 | 94.70 | 96.69 |
| Allowed (%) | 4.68 | 5.30 | 3.31 |
| Disallowed (%) | 0 | 0 | 0 |

Supplementary Table 2: Molecular dynamics simulation system information

| System | System shape | System size (Å³) | Number of components | | | | |
|---|---|---|---|---|---|---|---|
| | | | Lipids (POPC) | Water (TIP3P) | $Na^+$ | $Cl^-$ | Total atoms |
| Maprotiline-bound WT, inactive conformation | Rectangular | 180 x 180 x 204 | 770 | 150899 | 412 | 432 | 622973 |
| Diclofenac-bound WT, inactive conformation, ligand configuration 1 | Rectangular | 180 x 180 x 211 | 767 | 157317 | 431 | 446 | 641805 |
| Diclofenac-bound WT, inactive conformation, ligand configuration 2 | Rectangular | 180 x 180 x 211 | 766 | 157079 | 430 | 445 | 640955 |
| Diclofenac-bound WT, active conformation, ligand configuration 1 | Hexagonal | 180 x 180 x 141 | 633 | 81722 | 223 | 255 | 366714 |
| Diclofenac-bound WT, active conformation, ligand configuration 2 | Hexagonal | 180 x 180 x 141 | 633 | 81895 | 224 | 256 | 367235 |
| Diclofenac-bound L915A, inactive conformation, ligand configuration 1 | Rectangular | 180 x 180 x 211 | 767 | 157301 | 431 | 446 | 641721 |
| Diclofenac-bound L915A, inactive conformation, ligand configuration 2 | Rectangular | 180 x 180 x 211 | 766 | 157095 | 431 | 446 | 640969 |
| VIP1.409-bound WT, inactive conformation, ligand | Hexagonal | 180 x 180 x 140 | 649 | 80580 | 218 | 254 | 365442 |

| | | | | | | | |
|---|---|---|---|---|---|---|---|
| configuration 1 | | | | | | | |
| VIP1.409-bound WT, inactive conformation, ligand configuration 2 | Hexagonal | 180 x 180 x 140 | 648 | 80561 | 218 | 254 | 365251 |
| VIP1.409-bound WT, active conformation, ligand configuration 1 | Hexagonal | 180 x 180 x 141 | 633 | 81724 | 223 | 259 | 366740 |
| VIP1.409-bound WT, active conformation, ligand configuration 2 | Hexagonal | 180 x 180 x 141 | 633 | 81793 | 223 | 259 | 366947 |
| VIP1.409-bound L915A, inactive conformation, ligand configuration 1 | Hexagonal | 180 x 180 x 140 | 649 | 80554 | 219 | 255 | 365330 |
| VIP1.409-bound L915A, inactive conformation, ligand configuration 2 | Hexagonal | 180 x 180 x 140 | 648 | 80492 | 218 | 254 | 365008 |

### Chemical Synthesis and Characterization

#### General Chemistry Methods

Reagents purchased from Sigma-Aldrich Chemicals, Ambeed Inc, Chemscene Inc, were used without further purification. While performing synthesis, reaction mixtures were purified by silica gel flash chromatography on E. Merck 230–400 mesh silica gel 60 using a Teledyne ISCO CombiFlash Rf instrument with UV detection at 280 and 254 nm. RediSep Rf silica gel normal phase columns were used with a gradient range of 0–100% MeOH in DCM and EtOAc in hexane. Reported yields are isolated yields upon purification of each intermediate. Final clean (purity ≥95%, LC-MS Agilent 1100 Series LC/MSD) compounds were used for the study. NMR spectra were collected using Varian 400 MHz and 500 MHz NMR instrument at the NMR facility of Washington University School of Medicine in St. Louis collected via the Bruker Topspin Software (Bruker Topspin 3.5 pl 6). Chemical shifts are reported in parts per million (ppm) relative to residual solvent peaks at the nearest 0.01 for proton and 0.1 for carbon ($CDCl_3$ $^1H$: 7.26, $^{13}C$: 77.1; and $CD_3OD$ 1H: 3.31, 13C: 49.0, DMSO(d6) H: 2.50, 13C: 39.52). Peak multiplicity is reported as the NMR spectra were processed with MestreNova software14.2.0, namely s – singlet, d – doublet, t – triplet, q – quartet, m – multiplet for examples. Coupling constant (J) values are expressed in Hz. Mass spectra were obtained at the St. Louis College of Pharmacy using the Agilent 1100 Series LC/MSD by electrospray (ESI) ionization with a gradient elution program (Ascentis Express Peptide C18 column, acetonitrile/water 5/95/95/5, 5 min, 0.05% formic acid) and UV detection (214 nM/254 nM). High resolution mass spectra were obtained using a Bruker 10 T APEX -Qe FTICR-MS and the accurate masses are reported for the molecular ion $[M+Na]^+$. Detail experiments and characterization of the new compounds are included in the supporting information section.

#### Procedure and Synthetic Schemes

**Methyl 2-(2-((2,6-dichlorophenyl)amino)phenyl)acetate (VIP4.109):** Diclofenac (0.1 g, 0.33 mmol) was dissolved in methanol (8 mL), and concentrated $H_2SO_4$ (0.3 mL) was added. The reaction mixture was heated at 65 °C for 15 min. After completion, the reaction mixture was allowed to cool to room temperature and then poured into ice water. The resulting suspension was kept for 30 min to facilitate precipitation. The precipitated solid was collected by filtration and purified by flash column chromatography using 5-10% ethyl acetate in hexane as the eluent to afford **(VIP4.109)** (71 mg, 64%). NMR is in accordance with the known literature[1]. $^1H$ NMR (500

MHz, $CDCl_3$) δ 7.36 (d, *J* = 8.1 Hz, 2H), 7.14 (t, *J* = 7.7 Hz, 1H), 7.03 – 6.93 (m, 4H), 6.56 (d, *J* = 8.0 Hz, 1H), 3.83 (s, 2H), 3.77 (s, 3H). HRMS calcd for $C_{15}H_{13}Cl_2NO_2\ Na^+$ 332.0215; found, 332.02160.

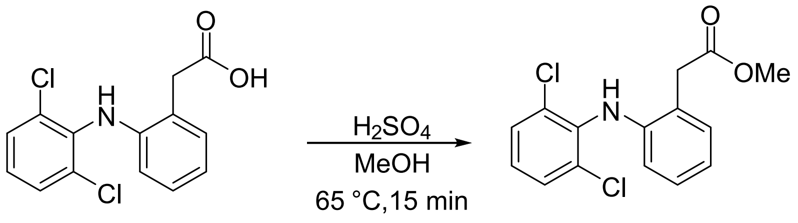


**VIP 4.109**

**2-(2-(phenylamino)phenyl)acetic acid:** It was purchased from Chem scene Inc. and used as such.

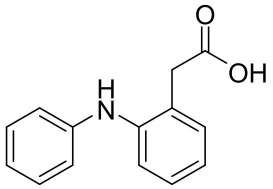


**2-[(2-Bromo-6-chlorophenyl)amino]phenylacetic acid:** It was purchased from Pharmaffiliates and used as such.

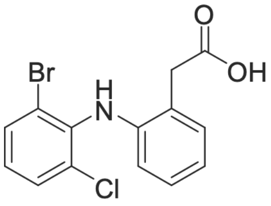


**2-(2-((2,6-difluorophenyl)amino)phenyl)acetic acid (VIP5.12):** Methyl 2-(2-iodophenyl)acetate (1.0 g, 3.26 mmol) and 2,6-difluoroaniline (0.701 g, 5.43 mmol) were dissolved in dry toluene (10 mL) under an argon atmosphere. $Pd_2(dba)_3$ (0.149 g, 0.16 mmol), rac-BINAP (0.125 g, 0.20 mmol), and $Cs_2CO_3$ (1.6 g, 4.89 mmol) were added sequentially, and the reaction mixture was heated at 110 °C for 12 h. After completion of the reaction, the solvent was removed under reduced pressure. The resulting residue was dissolved in water (10 mL) and extracted with ethyl acetate (2 × 15 mL). The combined organic extracts were washed with brine, dried over anhydrous sodium sulfate, filtered, and concentrated under reduced pressure. The crude product was purified by flash column chromatography using 5-10% ethyl acetate in hexane as the eluent. The purified ester intermediate was then subjected to hydrolysis using 1 N aqueous NaOH (2 mL) in ethanol (8 mL). The reaction mixture was refluxed at 77 °C for 12 h. After completion, the solvent was removed under reduced pressure, and the residue was diluted with water and acidified with 2 N HCl. The resulting mixture was extracted with ethyl acetate (2 × 15 mL). The combined organic layers were washed with brine, dried over anhydrous sodium sulfate, filtered, and concentrated under reduced pressure. The crude product was purified by flash column chromatography using 40-50% ethyl acetate in hexane to afford the desired product, HPLC purification (32 mg, 4%). $^1$H NMR (399 MHz, $CD_3OD$) δ 7.22 – 7.14 (m, 1H), 7.12 (d, $J$ = 8.2 Hz, 0H), 7.10 – 6.94 (m, 2H), 6.89 (t, $J$ = 6.8 Hz, 0H), 6.66 (d, $J$ = 7.9 Hz, 0H), 3.72 (s, 2H). $^{13}$C NMR (100 MHz, $CD_3OD$) δ 174.6, 157.3, 157.2, 154.9, 154.8, 142.6, 130.3, 127.5, 124.1, 122.0, 121.0, 116.3, 111.5, 111.46, 111.4, 111.3, 37.7. HRMS calcd for $C_{14}H_{11}F_2NO_2$ $Na^+$ 286.0650; found, 286.0650.

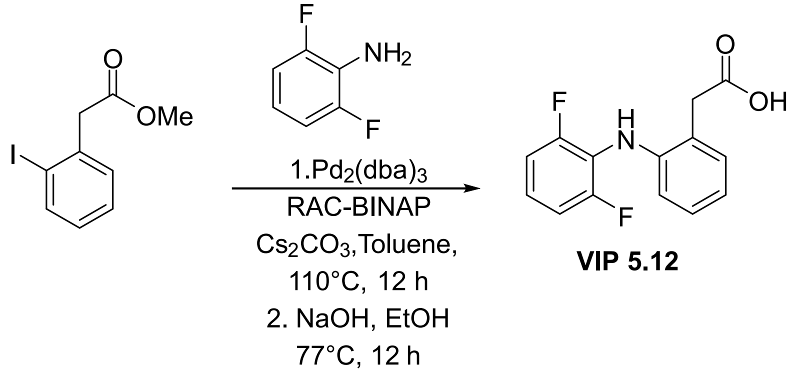


**VIP 5.12**

**2-(2-((2-chloro-6-fluorophenyl)amino)phenyl) acetic acid (VIP5.13)**: Following a known protoco[2] 2-(2-iodophenyl) acetic acid (0.2 g, 0.76 mmol) and 2-chloro6-fluoroaniline (0.44 g, 3.05 mmol) were dissolved in NMP (5 mL), to this CuI (0.288 g, 1.52 mmol) and $K_2CO_3$ (528 mg, 3.8 mmol) were added sequentially and reaction mixture was heated at 100 °C for 12 h. Reaction mixture cooled at rt and diluted with EtOAc (20 mL), water (5 mL), Conc. HCl (0.5 mL), charcoal (1 g), and celite ( 2 g) were added sequentially and stirred for 1 h. Mixture was filtered to give two layers, organic phase separated and dried over anhydrous sodium sulfate, filtered, and concentrated under reduced pressure. The crude product was purified by flash column chromatography using 30-40% ethyl acetate in hexane to afford the desired product which was subjected for HPLC purification (7 mg, 3%). 1H NMR (399 MHz, cdcl3) δ 7.26 (d, J = 1.5 Hz, 2H),

7.26 – 7.13 (m, 1H), 7.02 – 6.90 (m, 3H), 6.78 – 6.70 (m, 2H), 3.83 (s, 3H). $^{13}C$ NMR (100 MHz, cdcl₃) δ 174.00, 130.90, 128.31, 125.54, 122.16, 117.99, 115.03, 114.83, 38.13. HRMS calcd for C14H11ClFNO2 Na+ = 302.0354; observed mass = 302.0355.

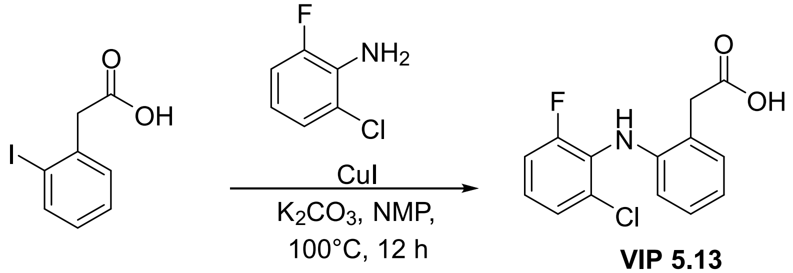


**VIP 5.13**

**2-(2-((2-chlorophenyl) amino) phenyl) acetic acid (VIP5.14):** 2-(2-iodophenyl) acetic acid (1 g, 3.81 mmol) and 2-chloroaniline (1.94 g, 15.26 mmol) were dissolved in NMP (8 mL), to this CuI (1.44 g, 7.62 mmol) and $K_2CO_3$ (2.64 g, 19.5) were added sequentially and reaction mixture was heated at 100 °C for 12 h. Reaction mixture cooled at rt and diluted with EtOAc (20 mL), water (5 mL), Conc. HCl (0.5 mL), charcoal (1 g), and celite (2 g) were added sequentially and stirred for 1 h. Mixture was filtered to give two layers, organic phase separated and dried over anhydrous sodium sulfate, filtered, and concentrated under reduced pressure. The crude product was purified by flash column chromatography using 30-40% ethyl acetate in hexane to afford the desired product which was subjected for HPLC purification (30 mg, 3%). $^{1}H$ NMR (399 MHz, $CDCl_3$) δ 7.39 – 7.24 (m, 4H), 7.14 – 7.02 (m, 2H), 6.96 (dd, *J* = 8.2, 1.6 Hz, 1H), 6.80 – 6.68 (m, 1H), 3.71 (s, 2H). $^{13}C$ NMR (100 MHz, $CDCl_3$) δ 177.8, 141.1, 140.8, 131.5, 129.8, 128.7, 127.3, 127.2, 124.07, 123.0, 121.2, 119.9, 115.3, 38.0.; HRMS calcd for $C_{14}H_{12}ClNO_2$ $Na^+$ 284.0448 observed mass 284.0449.

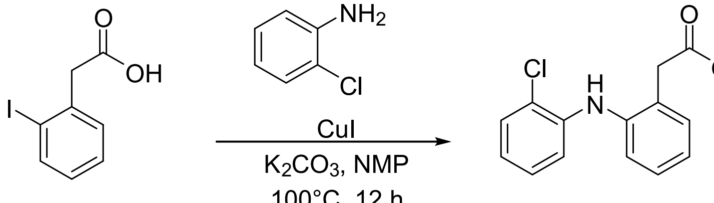


**VIP 5.14**

**2-(2-((2-bromo-6-fluorophenyl)amino)phenyl)acetic acid (VIP 5.18)**: 2-(2-Iodophenyl)acetic acid (1.0 g, 3.81 mmol) and 2-bromo-6-fluoroaniline (2.89 g, 15.26 mmol) were dissolved in DMF (8 mL). CuI (1.44 g, 7.62 mmol) and $K_2CO_3$ (2.64 g, 19.5 mmol) were added sequentially, and the reaction mixture was heated at 100 °C in a microwave reactor for 1 h. After completion, the reaction mixture was cooled to room temperature and diluted with EtOAc (20 mL) and water (5 mL). concentrated HCl (0.5 mL), activated charcoal (1 g), and Celite® (2 g) were added sequentially, and the resulting mixture was stirred for 1 h. The mixture was then filtered, and the layers were separated. The organic layer was dried over anhydrous sodium sulfate, filtered, and concentrated under reduced pressure. The crude product was purified by flash column chromatography using 30-40% ethyl acetate in hexane as the eluent to afford the desired product, which was further purified by preparative HPLC to give the title compound as the final product (53 mg, 4% yield). $^1$H NMR (399 MHz, DMSO d6) δ 7.51 (d, *J* = 8.1 Hz, 0H), 7.28 (t, *J* = 9.8 Hz, 1H), 7.06 (t, *J* = 8.2 Hz, 1H), 6.85 (d, *J* = 7.9 Hz, 1H), 6.42 (d, *J* = 8.0 Hz, 1H), 3.66 (s, 2H). $^{13}$C NMR (100 MHz, DMSO d6) δ 173.8, 157.8, 155.4, 143.2, 131.3, 130.33, 129.4, 129.37, 128.0, 125.5, 125.5, 124.3, 121.2, 119.5, 116.5, 116.4, 116.3, 110.0, 38.3. HRMS calcd for $C_{14}H_{11}BrFNO_2Na^+$ 345.9849 observed mass 345.9849.

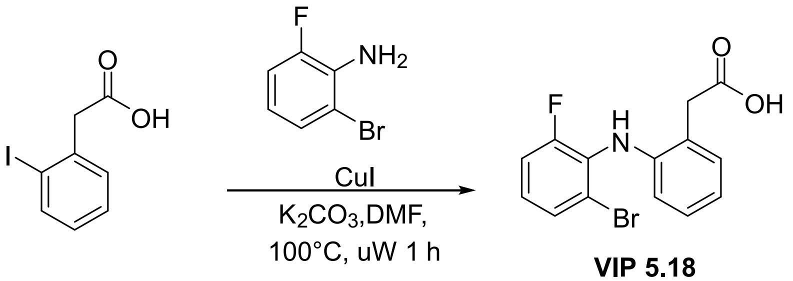


**VIP 5.18**

**2-(2-((2-chloro-6-iodophenyl)amino)phenyl)acetic acid (VIP 5.19):** Using a procedure similar to **VIP 5.18**, 2-(2-iodophenyl)acetic acid (1.0 g, 3.81 mmol) and 2-chloro-6-iodoaniline (3.85 g, 15.26 mmol) were dissolved in DMF (8 mL). CuI (1.44 g, 7.62 mmol) and $K_2CO_3$ (2.64 g, 19.5 mmol) were added sequentially, and the reaction mixture was heated at 100 °C in a microwave reactor for 1 h. After workup and purification as described for VIP 5.18, the desired product was obtained (21 mg, 2% yield). $^1$H NMR (399 MHz, DMSO d6) δ 7.87 (d, *J* = 7.8 Hz, 1H), 7.55 (d, *J* = 8.0 Hz, 1H), 7.18 (d, *J* = 7.5 Hz, 1H), 7.10 (s, 1H), 7.02 (t, *J* = 7.7 Hz, 1H), 6.95 (t, *J* = 7.9 Hz, 1H), 6.82 (t, *J* = 7.4 Hz, 1H), 6.16 (d, *J* = 8.0 Hz, 1H), 3.67 (s, 2H). $^{13}$C NMR (100 MHz, DMSO d6) δ 173.7, 143.6, 141.6, 139.2, 131.3, 130.9, 130.2, 128.00, 127.89, 124.21, 120.8, 116.1, 99.7, 38.3. HRMS calcd for $C_{14}H_{11}ClINO_2$ $Na^+$ 409.941524, observed mass 409.94159.

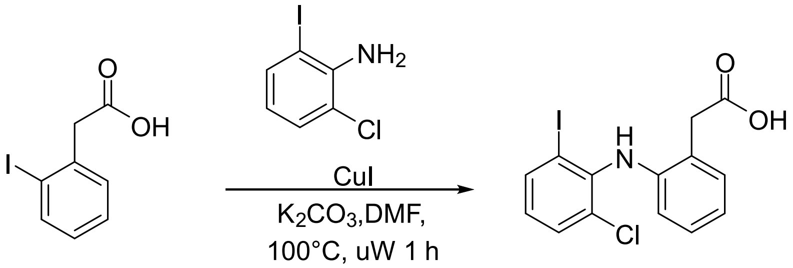


**VIP 5.19**

**2-(2-((2-chloro-6-methylphenyl)amino)phenyl)acetic acid (VIP 5.20):** Using a procedure similar to VIP 5.18, 2-(2-iodophenyl)acetic acid (1.0 g, 3.81 mmol) and 2-chloro-6-methylaniline (2.15 g, 15.26 mmol) were dissolved in DMF (8 mL). CuI (1.44 g, 7.62 mmol) and $K_2CO_3$ (2.64 g, 19.5 mmol) were added sequentially, and the reaction mixture was heated at 100 °C in a microwave reactor for 1 h. After workup and purification as described for VIP 5.18, the desired product was obtained (62 mg, 6% yield). $^1$H NMR (399 MHz, DMSO d6) δ 12.55 (s, 1H), 7.36 (d, *J* = 8.0 Hz, 1H), 7.22 (d, *J* = 7.5 Hz, 1H), 7.14 – 7.04 (m, 1H), 7.04 – 6.95 (m, 1H), 6.92 (s, 1H), 6.75 (t, *J* = 7.0 Hz, 1H), 6.11 (d, *J* = 7.5 Hz, 1H), 3.69 (s, 2H), 2.04 (s, 3H). $^{13}$C NMR (100 MHz, DMSO d6) δ 173.7, 144.1, 138.2, 136.4, 131.4, 130.4, 130.2, 128.1, 127.9, 125.8, 123.1, 119.9, 114.7, 38.1, 18.9. HRMS calcd for $C_{15}H_{14}ClNO_2$ $Na^+$ 298.0605, observed mass 298.0605.

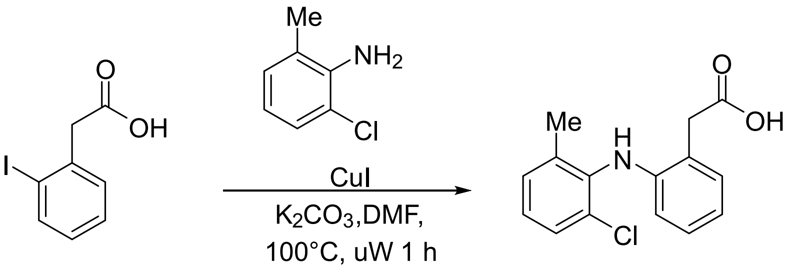


**VIP 5.20**

**2-(2-((2-fluoro-6-iodophenyl)amino)phenyl)acetic acid (VIP5.24):** Using a procedure similar to VIP 5.18, 2-(2-iodophenyl)acetic acid (1.0 g, 3.81 mmol) and 2-fluoro-6-iodoaniline (3.61 g, 15.26

mmol) were dissolved in DMF (8 mL). CuI (1.44 g, 7.62 mmol) and $K_2CO_3$ (2.64 g, 19.5 mmol) were added sequentially, and the reaction mixture was heated at 100 °C in a microwave reactor for 1 h. After workup and purification as described for VIP 5.18, the desired product was obtained (25 mg, 2% yield). $^1$H NMR (399 MHz, DMSO d6) δ 7.71 (d, *J* = 8.0 Hz, 1H), 7.33 – 7.23 (m, 1H), 7.17 (d, *J* = 6.0 Hz, 1H), 7.03 (t, *J* = 7.7 Hz, 1H), 6.99 – 6.94 (m, 2H), 6.81 (t, *J* = 7.4 Hz, 1H), 6.32 (d, *J* = 8.0 Hz, 1H), 3.66 (s, 2H). $^{13}$C NMR (100 MHz, DMSO d6) δ 173.66, 157.63, 155.15, 143.61, 135.60, 135.56, 133.03, 132.90, 131.24, 127.98, 127.17, 127.09, 123.86, 120.76, 117.14, 116.93, 115.98, 98.36, 38.14. HRMS calcd for $C_{14}H_{11}FINO_2Na^+$ 393.9710, observed mass = 393.9711.

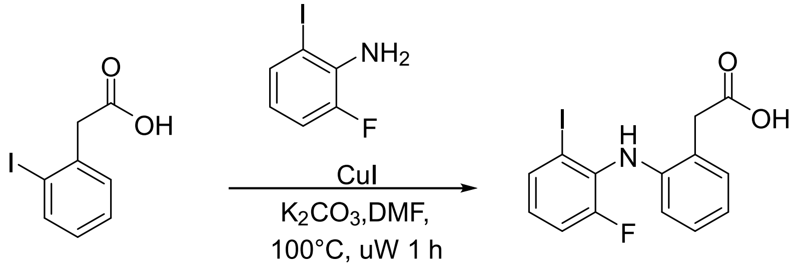


**VIP 5.24**

**2-(2-((2,6-dimethylphenyl)amino)phenyl)acetic acid (VIP5.33)**: Using a procedure similar to VIP 5.18, 2-(2-iodophenyl)acetic acid (1.0 g, 3.81 mmol) and 2,6-dimethylaniline (1.86 g, 15.26 mmol) were dissolved in DMF (8 mL). CuI (1.44 g, 7.62 mmol) and $K_2CO_3$ (2.64 g, 19.5 mmol) were added sequentially, and the reaction mixture was heated at 100 °C in a microwave reactor for 1 h. After workup and purification as described for VIP 5.18, the desired product was obtained (20 mg, 2% yield). $^1$H NMR (399 MHz, $CDCl_3$) δ 7.20 (d, *J* = 7.9 Hz, 1H), 7.14 – 7.00 (m, 5H), 6.80 (t, *J* = 7.4 Hz, 2H), 6.29 (d, *J* = 8.1 Hz, 1H), 3.79 (s, 2H), 2.13 (s, 6H). $^{13}$C NMR (100 MHz, $CDCl_3$) δ 174.7, 144.7, 138.9, 134.0, 131.1, 128.7, 128.6, 124.8, 120.16, 119.3, 114.7, 38.3, 18.3. HRMS calcd for $C_{16}H_{17}NO_2$ $Na^+$ 278.1151 observed mass 278.1151.

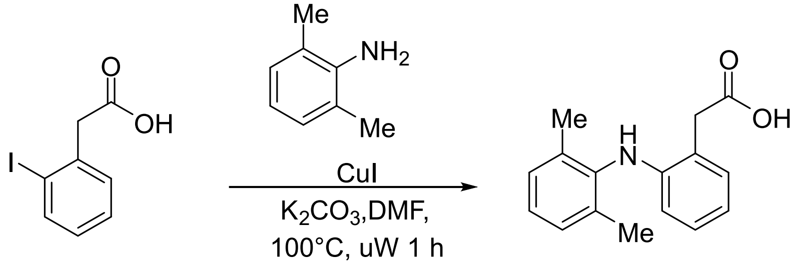


**VIP 5.33**

**2-(2-((2,6-dichlorophenyl)amino)phenyl)-N,N-dimethylacetamide: (VIP5.37)**: To the diclofenac (0.2 g, 0.67 mmol) dissolved in dichloromethane (10 mL) was added dimethyl amine (2N in THF, 6.75 ml, 13.51 mmol) and stirred for a minute. HATU (0.51 g, 1.34 mmol), DIPEA (0.35 mL, 2.01 mmol) were added sequentially and reaction mixture stirred for 24 h. Reaction mixture diluted with EtOAc (20 mL) and washed with water and brine. Organic phase separated and dried over anhydrous sodium sulfate, filtered, and concentrated under reduced pressure. The crude product was purified by flash column chromatography using 40-60% ethyl acetate in hexane (0.11 g, 58%). $^{1}$H NMR (399 MHz, $CDCl_3$) δ 7.79 (s, 1H), 7.33 (d, *J* = 8.1 Hz, 2H), 7.16 (d, *J* = 6.7 Hz, 1H), 7.09 (td, *J* = 7.7, 1.6 Hz, 1H), 6.96 (t, *J* = 8.0 Hz, 1H), 6.89 (td, *J* = 7.4, 1.3 Hz, 1H), 6.51 (d, *J* = 8.0 Hz, 1H), 3.85 (s, 2H), 3.21 (s, 3H), 3.00 (s, 3H). $^{13}$C NMR (100 MHz, $CDCl_3$) δ 171.6, 143.7, 138.0, 130.5, 130.1, 128.8, 127.5, 124.8, 123.9, 121.0, 117.5, 77.4, 77.0, 76.7, 38.1, 37.6, 36.0. HRMS calcd for $C_{16}H_{16}Cl_2N_2O$ $Na^+$ 345.0531 observed mass = 345.0532

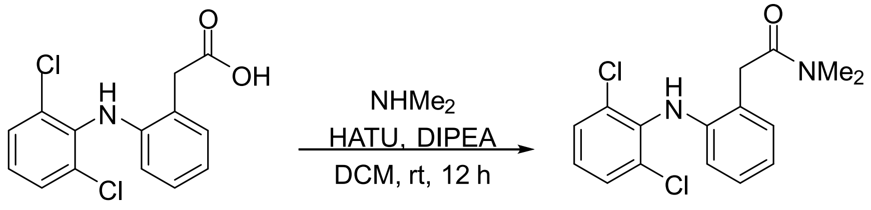


**VIP 5.37**

**2-(2-((2,6-dichlorophenyl)amino)phenyl)-N-methylacetamide** (**VIP5.52**): To the diclofenac (0.5 g, 1.68 mmol) dissolved in dichloromethane (10 mL) was added methyl amine (2N in 40% in $H_2O$, 2.55 ml, 33 mmol) and stirred for a minute. HATU (0.957 g, 2.52 mmol), DIPEA (0.43 mL, 2.52 mmol) were added sequentially and reaction mixture stirred for 24 h. Reaction mixture diluted with EtOAc (20 mL) and washed with water and brine. Organic phase separated and dried over anhydrous sodium sulfate, filtered, and concentrated under reduced pressure. The crude product was purified by flash column chromatography using 60% ethyl acetate in hexane (0.135 g, 26%). $^{1}$H NMR (399 MHz, $CDCl_3$) δ 7.41 – 7.31 (m, 1H), 7.18 – 7.06 (m, 2H), 6.98 (t, *J* = 8.0 Hz, 2H), 6.94 – 6.85 (m, 1H), 6.51 (d, *J* = 8.0 Hz, 1H), 6.26 (q, *J* = 4.9 Hz, 1H), 3.67 (s, 2H), 2.78 (d, *J* = 4.9 Hz, 3H). $^{13}$C NMR (100 MHz, $cdcl_3$) δ 172.4, 142.9, 137.5, 130.7, 130.0, 128.9, 127.9, 124.6, 124.3, 121.5, 117.4, 40.8, 26.6. LCMS calcd for $C_{15}H_{14}Cl_2N_2O+H^+$ 308.04 observed mass = 309.2

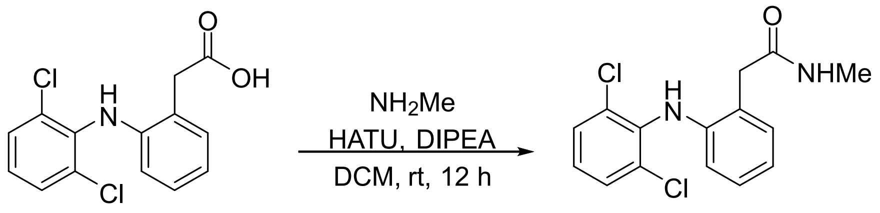

**VIP 5.52**

**2-(2-((2,6-dichlorophenyl)amino)phenyl)acetamide** (**VIP5.53**): To the diclofenac (0.5 g, 1.68 mmol) dissolved in dichloromethane (10 mL) was added $NH_4OH$ (28% in $H_2O$, 4.13 ml, 33 mmol) and stirred for a minute. HATU (0.957 g, 2.52 mmol), DIPEA (0.43 mL, 2.52 mmol) were added sequentially and reaction mixture stirred for 24 h. Reaction mixture diluted with EtOAc (20 mL) and washed with water and brine. Organic phase separated and dried over anhydrous sodium sulfate, filtered, and concentrated under reduced pressure. The crude product was purified by flash column chromatography using 60% ethyl acetate in hexane (140 mg, 29%). $^1$H NMR (399 MHz, DMSO (d6)) δ 8.45 (s, 1H), 7.80 (s, 1H), 7.49 (d, *J* = 8.1 Hz, 2H), 7.16 (dd, *J* = 16.3, 7.6 Hz, 1H), 7.02 (t, *J* = 7.7 Hz, 1H), 6.83 (t, *J* = 7.2 Hz, 1H), 6.27 (d, *J* = 8.0 Hz, 1H), 3.52 (s, 2H). $^{13}$C NMR (100 MHz, DMSO (d6)) δ 174.4, 143.5, 143.3, 137.7, 130.9, 129.82, 129.8, 129.7, 127.6, 125.9, 125.5, 121.10, 121.05, 116.3, 116.3, 39.4. LCMS calcd for $C_{14}H_{12}Cl_2N_2O+H^+$ 294.03 observed mass = 295.1.

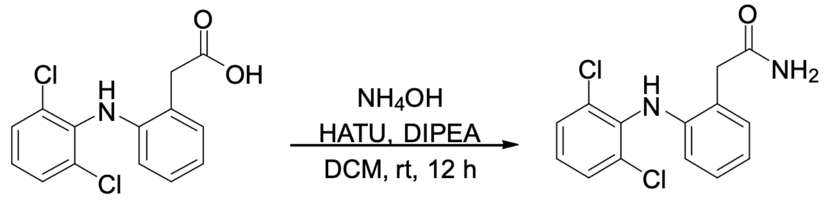

**VIP 5.53**

**Ethyl 2-(2-((2,6-dichlorophenyl)amino)phenyl)acetate (VIP 5.54**): Using a similar procedure to VIP4.109**,** diclofenac (0.3 g, 1.01 mmol) was dissolved in ethanol (8 mL), and concentrated $H_2SO_4$ (0.3 mL) was added. The reaction mixture was heated at 78 °C for 15 min. After workup and purification as described for VIP 4.109, the desired product was obtained (266 mg, 81% yield).

$^1H$ NMR (399 MHz, $CDCl_3$) δ 7.35 (d, *J* = 8.1 Hz, 2H), 7.24 (d, *J* = 7.6 Hz, 1H), 7.17 – 7.09 (m, 1H), 7.03 – 6.92 (m, 2H), 6.56 (d, *J* = 8.0 Hz, 1H), 4.26 – 4.15 (m, 2H), 3.81 (s, 2H), 1.30 (t, *J* = 7.1 Hz, 3H). $^{13}C$ NMR (100 MHz, $CDCl_3$) δ 172.4, 142.7, 137.9, 130.9, 129.5, 128.9, 127.9, 124.4, 124.0, 122.0, 118.2, 77.3, 77.0, 76.7, 61.4, 38.8, 14.2. HRMS calcd for $C_{16}H_{15}Cl_2NO_2Na^+$ 346.0372 observed mass 346.0372.

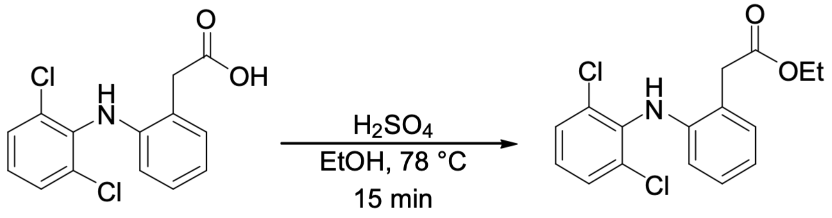


**VIP 5.54**

**Isopropyl 2-(2-((2,6-dichlorophenyl)amino)phenyl)acetate (VIP5.55):** Using a similar procedure to VIP4.109**,** diclofenac (0.2g, 1.01 mmol) was dissolved in ethanol (8 mL), and concentrated $H_2SO_4$ (0.3 mL) was added. The reaction mixture was heated at 78 °C for 15 min. After workup and purification as described for VIP 4.109, the desired product was obtained (170 mg, 78% yield). $^1H$ NMR (399 MHz, $CDCl_3$) δ 7.34 (d, *J* = 8.0 Hz, 2H), 7.28 – 7.20 (m, 1H), 7.12 (td, *J* = 8.1, 1.6 Hz, 1H), 7.02 – 6.91 (m, 3H), 6.56 (d, *J* = 8.0 Hz, 1H), 5.07 (p, *J* = 6.3 Hz, 1H), 3.78 (s, 2H), 1.27 (d, *J* = 6.3 Hz, 6H). $^{13}C$ NMR (100 MHz, $CDCl_3$) δ 172.0, 142.8, 138.0, 130.8, 129.5, 128.8, 127.8, 124.8, 123.9, 122.0, 118.4, 77.3, 77.0, 76.7, 68.7, 39.0, 21.8. HRMS calcd for $C_{17}H_{17}Cl_2NO_2$ $Na^+$ 360.0528 observed mass 360.0529.

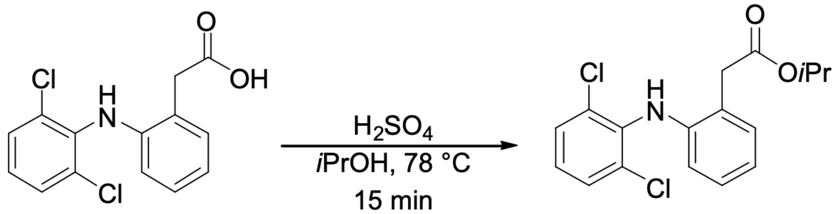


**VIP 5.55**

**Methyl 2-(2-((2-chloro-6-iodophenyl)amino)phenyl)acetate (VIP5.67):** Using a similar procedure to VIP4.109**,** VIP5.19 (15 mg, 0.038 mmol) was dissolved in methanol (5 mL), and concentrated $H_2SO_4$ (0.05 mL) was added. The reaction mixture was heated at 65 °C for 15 min.

After workup and purification as described for VIP 4.109, the desired product was obtained (10 mg, 65% yield). $^{1}$H NMR (500 MHz, $CDCl_3$) δ 7.79 (dd, *J* = 7.9, 1.4 Hz, 1H), 7.44 (dd, *J* = 8.0, 1.4 Hz, 1H), 7.29 – 7.24 (m, 1H), 7.14 (dd, *J* = 7.7, 1.6 Hz, 1H), 7.00 – 6.94 (m, 1H), 6.84 – 6.77 (m, 2H), 6.48 (d, *J* = 8.0 Hz, 1H), 3.84 (s, 2H), 3.77 (s, 3H). $^{13}$C NMR (126 MHz, $CDCl_3$) δ 172.6, 143.0, 141.7, 138.7, 131.0, 130.5, 129.3, 128.2, 126.0, 124.3, 121.9, 118.1, 96.3, 52.5, 38.6. HRMS calcd for $C_{15}H_{13}ClINO_2$ $Na^+$ 423.9571, observed mass 423.9572.

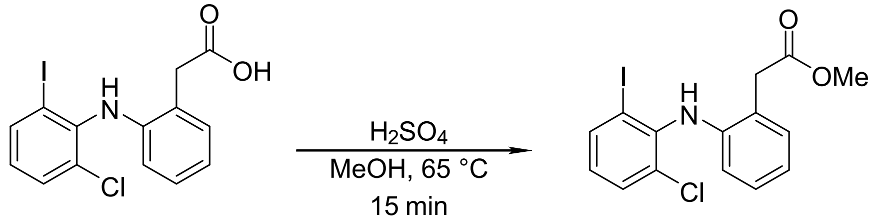


**VIP 5.19** **VIP 5.67**

<u>References</u>


1. Majhi et. al. Regioselective Iodination of Arenes Using Iron- or Silver-Catalyzed Activation of N-Iodosaccharin. *J. Org. Chem*. 2026, 91, 2257−2267.
2. Mei et. al. Expedient Drug Synthesis and Diversification via ortho-C−H Iodination using Recyclable PdI2 as the Precatalyst. *Org. Lett.* 2010, 12, 3140–3143
